# Dual Mechanisms for Heterogeneous Responses of Inspiratory Neurons to Noradrenergic Modulation

**DOI:** 10.1101/2025.07.25.666713

**Authors:** Sreshta Venkatakrishnan, Andrew K. Tryba, Alfredo J. Garcia, Yangyang Wang

## Abstract

Respiration is an essential involuntary function necessary for survival. This poses a challenge for the control of breathing. The preBötzinger complex (preBötC) is a heterogeneous neuronal network responsible for driving the inspiratory rhythm. While neuromodulators such as norepinephrine (NE) allow it to be both robust and flexible for all living beings to interact with their environment, the basis for how neuromodulation impacts neuron-specific properties remains poorly understood. In this work, we examine how NE influences different preBötC neuronal subtypes by modeling its effects through modulating two key parameters: calcium-activated nonspecific cationic current gating conductance (*g*_CAN_) and inositol-triphosphate (IP_3_), guided by experimental studies. Our computational model captures the experimentally observed differential effects of NE on distinct preBötC bursting patterns. We show that this dual mechanism is critical for inducing conditional bursting and identify specific parameter regimes where silent neurons remain inactive in the presence of NE. Furthermore, using methods of dynamical systems theory, we uncover the mechanisms by which NE differentially modulates burst frequency and duration in NaP-dependent and CAN-dependent bursting neurons. These results align well with previously reported experimental findings and provide a deeper understanding of cell-specific neuromodulatory responses within the respiratory network.

**MSC codes:** 37N25, 34C23, 34C60, 34E13, 34E15, 92C20

## 1. Introduction

Breathing is a complex neurophysiological process required for survival and consists of two main phases — inspiration and expiration. In mammals, the preBötzinger complex (preBötC) within the brainstem generates the neural rhythm that drives inspiration and is involved with regulating the phasic timing of inspiration and expiration. Rhythmic activity persists in some preBötC neurons even when synaptic coupling in the network is blocked and the network rhythm ceases. Such persistent activity is derived from intrinsic membrane properties among these so-called “pacemaker neurons” [49, 32, 23, 44]. Pacemaker neurons in the preBötC exhibit spontaneous rhythmic bursting activity, where a discrete barrage of action potentials is generated in a periodic manner, and are hence also referred to as intrinsic bursting neurons or bursters. Multiple mechanisms underlie bursting, including the calcium-activated non-specific cationic current (*I*_CAN_), driven by calcium oscillations within the preBötC, and the voltage-sensitive persistent sodium current (*I*_NaP_) [14, 33, 40, 39, 44, 37].

Experimental observations have inspired numerous computational studies (e.g., [9, 10, 47, 31, 51, 38]), aimed at elucidating the cellular mechanisms underlying intrinsic rhythmic bursting activity in individual preBötC neurons, the dynamic interplay between intrinsic bursting conductances, and their impact at the network level. An early influential computational model of isolated preBötC neurons only captures bursting driven by the *I*_NaP_-dependent mechanism [9]. Later models incorporated synaptic activation of *I*_CAN_ [47], followed by studies that combined both *I*_NaP_- and *I*_CAN_-dependent bursting mechanisms (e.g., [31, 51, 38, 43]). Beyond computational modeling, dynamical systems theory has provided valuable insights into the interplay between these two bursting mechanisms, revealing complex mixed bursting dynamics characterized by distinct bursting patterns [57, 59]. Notably, while some of these mixed bursting patterns exhibit significant qualitative similarities, mathematical analysis suggests they arise from fundamentally different mechanisms [58], underscoring the importance of complementing computational models with dynamical systems analysis.

Since the activity of these neurons is also constantly modulated by various neuromodulators via altering the intrinsic properties of neurons as well as properties of the network, several works have been dedicated to studying the effects of neuromodulation on the preBötC neurons [19, 18, 51, 60, 56]. Various neuromodulators such as serotonin, norepinephrine, acetylcholine, substance P have been shown to increase the frequency of the respiratory rhythmic activity [19, 2, 56, 48, 45, 26, 4, 54]. These neuromodulators can have unique effects on the different types of preBötC neurons, altering neuronal excitability and regularity of activity patterns, which, in turn, can enhance or diminish the stability of the network rhythm. Indeed, neuromodulation of the preBötC by norepinephrine (NE) stimulates rhythmogenesis and differentially affects activity patterns of synaptically isolated preBötC neurons [56]. Specifically, NE stimulated the burst frequency without affecting burst duration in *I*_NaP_-dependent bursting neurons. Whereas, NE increased burst duration of *I*_CAN_-dependent bursters while minimally affecting their burst frequency. Moreover, NE had differential effects on different synaptically isolated non-bursting neurons. Silent non-bursting neurons (i.e., neurons that did not generate action potentials in synaptic isolation) remained silent in NE. Whereas, in active non-bursting neurons (i.e., neurons continued to generate action potentials but did not burst in synaptic isolation), NE induced conditional bursting properties that are *I*_CAN_-dependent [56].

Previously, computational works such as [51, 38] have attempted to model the effects of NE on single preBötC neurons via increasing the conductance for the CAN-current (*g*_CAN_). Their choice in modeling the application of NE through the model parameter *g*_CAN_ was attributed to earlier findings that NE, which is an *α*_1_-receptor agonist, increases CAN-current conductances in various cell types. Both models successfully reproduced the effects of NE on synaptically isolated intrinsic bursting preBötC neurons as observed in *in vitro* experiments in [56] in terms of the frequency and duration of the bursts (see subsection 2.3 for details). However, increasing *g*_CAN_ in these *in silico* models did not produce the conditional bursting properties that were observed experimentally, nor did it capture the NE-insensitivity of silent non-bursters, suggesting that additional mechanisms are needed to comprehensively capture the experimental effects of NE on single preBötC neurons [56]. This paper addresses this gap by integrating additional mechanisms through with NE modulates neuronal activity. We hypothesize that NE enhances second messenger-mediated Ca^2+^ flux, a mechanism previously described in [21, 13]. Our modeling reveals that interactions between *g*_CAN_ and inositol-triphosphate (IP_3_) are critical for inducing conditional bursting and identifies discrete parameter regimes where silent non-bursting neurons continue to remain inactive even in the presence of NE. Furthermore, our model simulations and analyses (detailed in section 3) uncover the mechanisms by which NE differentially modulates burst frequency and duration in NaP-dependent and CAN-dependent bursting neurons. These findings align closely with experimental data, supporting the model’s relevance for probing the mechanisms underlying NE modulation of preBötC neurons.

The paper is organized as follows: subsection 2.1 presents a mathematical model of a single preBötC neuron that builds on previous models with modifications to facilitate the modeling of NE effects. subsection 2.2 outlines the various activity patterns observed in this model. subsection 2.3 introduces our proposed mechanism for NE modulation of preBötC neurons. In subsection 2.4, we review the mathematical tools used for analyzing the model. Section 3 presents our main results and detailed analysis of NE’s effects on NaP-dependent bursters (subsection 3.1), CAN-dependent bursters (subsection 3.2), Tonic Spiking neurons (subsection 3.3) and Quiescent neurons (subsection 3.4). Finally, we conclude with a discussion in section 4.

## 2. Mathematical Modeling and Preliminaries

In this section, we first present a single preBötC neuron model, followed by an outline of the various activity patterns it exhibits. We then introduce our proposed NE modulation mechanism and conclude by reviewing the geometric singular perturbation theory used in our analysis.

### 2.1. preBötC neuron Model

The preBötC neuron model considered in this paper is a single compartment dynamical system that incorporates Hodgkin-Huxley-style conductances, adapted from previously described models [51, 38, 59]. The model is described by the following equations:

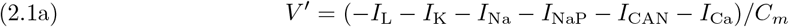

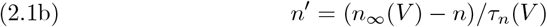

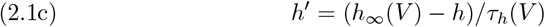

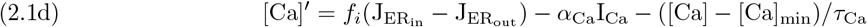

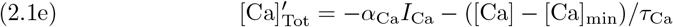

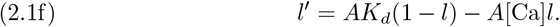

Equations (2.1a)-(2.1c), henceforth referred to as *the voltage subsystem*, describe the membrane potential dynamics *V* and the voltage-dependent activation *n* and inactivation *h* variables, respectively. The model includes a leak current (*I*_L_), a spike-generating fast potassium (*I*_K_) current, a fast spiking sodium (*I*_Na_) current, a persistent sodium current (*I*_NaP_), and a calcium-activated nonspecific cationic current (*I*_CAN_), through which calcium dynamics [Ca] influence membrane potential dynamics *V*. In addition to the currents from [51, 38], we incorporate an additional voltage-dependent calcium current [59, 42, 43] to account for the bidirectional interaction between membrane potential and calcium dynamics, which plays an important role in capturing the effects of NE, as discussed later in subsection 2.3. As a result, the neuron should no longer be considered closed to calcium flux into and out of the cell, as in [51, 38], where the total intracellular calcium concentration within the cell ([Ca]_Tot_) was assumed constant. Instead, model (2.1) now represents an open cell in which [Ca]_Tot_ is treated as a dynamic variable.

The ionic currents are defined as follows:

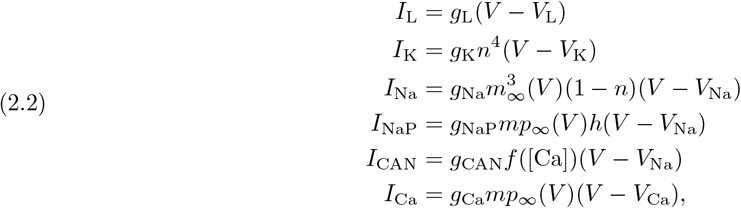

where *g*_*i*_ is the maximum conductance and *V*_*i*_ denotes the reversal potential for each current *I*_*i*_. To simplify the model, we followed prior studies [51, 38, 53, 11, 58] in our treatment of CAN and calcium currents, adopting similar formulations and parameter choices. The steady-state activation/inactivation (*x*_∞_) and time constant (*τ*_*x*_) for *I*_Na_, *I*_NaP_ and *I*_Ca_ take the following forms:

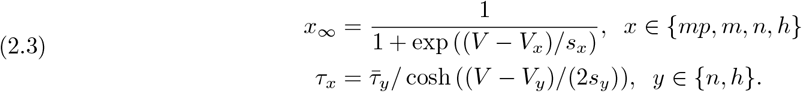

*I*_CAN_ activation takes a different form and depends on the calcium concentration in the cytoplasm [Ca]:

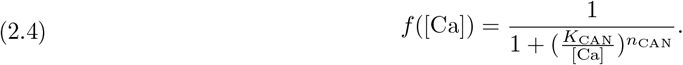

Equations (2.1d) - (2.1f), henceforth referred to as *the calcium subsystem*, describe the dynamics of intra-cellular calcium concentration in the cytoplasm [Ca], the combined intracellular and endoplasmic reticulum (ER) total calcium concentration [Ca]_Tot_, and the fraction of IP_3_ receptors that have not been inactivated. The calcium dynamics are influenced by the voltage-dynamics via the term − *α*_Ca_*I*_Ca_ in (2.1d) and (2.1e), which represents Ca^2+^ influx from the extracellular space through voltage-gated calcium channels. In (2.1d), 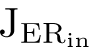 represents the calcium flux per unit volume from the endoplasmic reticulum (ER) into the cell cytoplasm, and this depends on *l*. 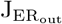 represents the calcium flux that flows out of the cytoplasm into the ER. These fluxes are modeled by (2.5a) - (2.5c):

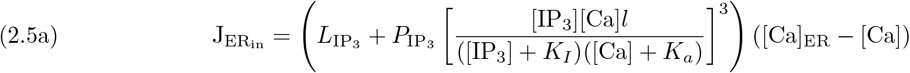

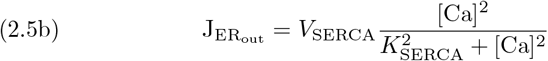

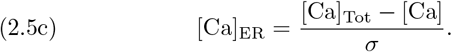

The last term in (2.1d) and (2.1e) represents the membrane Ca^2+^ pump, which expels free intracellular Ca^2+^ from the cytoplasm, where [Ca]_min_ sets a minimal baseline calcium concentration and *τ*_Ca_ is the time constant for the Ca^2+^ pump [31, 43, 47].

The parameter values associated with this model are listed in Table 1. Additional details about the model can be found in [38, 59, 31].

**Table 1:** The parameter values for model (2.1).

| Currents | Associated Parameter Values |
| --- | --- |
| $I_L$ | $g_L = 2.3\text{nS}$, $V_L = -58\text{mV}$. |
| $I_K$ | $g_K = 11.2\text{nS}$, $V_K = -85\text{mV}$. |
| $I_{\text{Na}}$ | $g_{\text{Na}} = 28\text{nS}$, $V_m = -34\text{mV}$, $V_n = -29\text{mV}$,<br>$s_m = -5\text{mV}$, $s_n = -4\text{mV}$, $\bar{\tau}_n = 10\text{ms}$, $V_{\text{Na}} = 50\text{mV}$. |
| $I_{\text{NaP}}$ | $g_{\text{NaP}} \in [0, 5]\text{nS}$, $V_h = -48\text{mV}$, $V_{mp} = -40\text{mV}$,<br>$s_h = 5\text{mV}$, $s_{mp} = -6\text{mV}$, $\bar{\tau}_h = 10000\text{ms}$. |
| $I_{\text{CAN}}$ | $g_{\text{CAN}} \in [0, 4]\text{nS}$, $K_{\text{CAN}} = 0.74\mu\text{M}$, $n_{\text{CAN}} = 0.97$. |
| $I_{\text{Ca}}$ | $g_{\text{Ca}} \in [0.00001, 0.0008]\text{nS}$, $V_{\text{Ca}} = 150\text{mV}$. |
| ER [Ca] | $[\text{IP}_3] \in [0, 1]\mu\text{M}$, $L_{\text{IP}_3} = 0.37/\text{ms}$, $P_{\text{IP}_3} = 31000/\text{ms}$,<br>$K_I = 1\mu\text{M}$, $K_a = 0.4\mu\text{M}$, $\sigma = 0.185$, $f_i = 0.000025$,<br>$V_{\text{SERCA}} = 400\mu\text{M}/\text{ms}$, $K_{\text{SERCA}} = 0.2\mu\text{M}$, $\alpha_{\text{Ca}} = 0.025\text{mM}/\text{fC}$,<br>$[\text{Ca}]_{\text{min}} = 0.005\mu\text{M}$, $\tau_{\text{Ca}} = 500\text{ms}$, $A = 0.005\mu\text{M}/\text{ms}$, $K_d = 0.4\mu\text{M}$. |
| Other | $C_m = 21\text{pF}$. |

### 2.2. Activity Patterns in the Model

The preBötC model (2.1) exhibits a diverse range of dynamically different activity patterns depending on variations in *g*_NaP_ and *g*_Ca_, the two critical parameters that govern transitions between different bursting mechanisms (see Figure 1(i)). The voltage and calcium time series of a dimensionless version of model (2.1) (see (2.6) in subsection 2.4) are shown for parameters corresponding to different regions in Figure 1(i). Specifically, panels illustrate (A) Quiescence, (B) *I*_NaP_-dependent bursting, (C) Tonic Spiking, (D) Mixed Bursting, and (E, F) two different types of *I*_CAN_-dependent bursting behaviors.

**Fig. 1:**
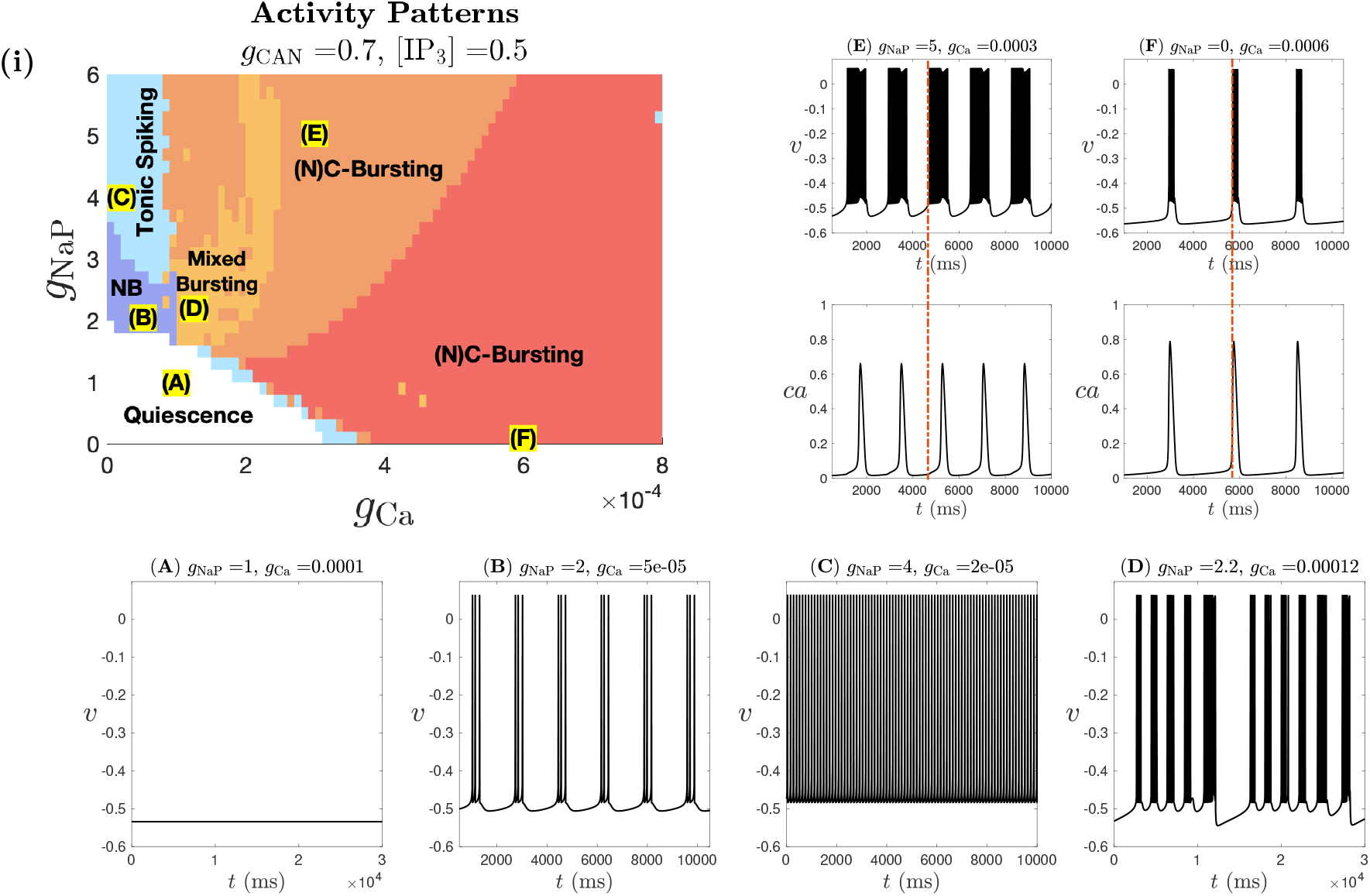
Activity patterns of the dimensionless model (2.6), derived from model (2.1), depend on parameter values *g*_NaP_ and *g*_Ca_, for *g*_CAN_ = 0.7 and [IP_3_] = 0.5. (i) The range of *g*_Ca_ and *g*_NaP_ for different solution patterns. White color denotes quiescence and other colors are tonic spiking (light blue) or bursting (dark blue, orange or red). Panels (A) through (F) illustrate sample traces for each activity pattern corresponding to labeled parameter values in panel (i). The red dashed line in panels (E) and (F) marks burst onset, highlighting a key difference between the two (N)C-Bursting regions (orange and red): in the orange region, the burst initiates before *ca* spikes, whereas in the red region, *ca* spikes before burst onset.

Increasing *g*_NaP_ at low *g*_Ca_ values causes transitions from quiescence (Figure 1(i), white region; Figure 1A) to NB (Figure 1(i), dark blue region; Figure 1B) and eventually to tonic spiking (Figure 1(i), pale blue region; Figure 1C). Bursting in the NB region occurs when the persistent sodium current is in the bursting range and there are no large Ca^2+^ oscillations. Following nomenclature established in prior studies [51, 38, 59], we refer to *I*_NaP_-dependent bursting as *N-bursting (NB)*. For relatively large *g*_Ca_ values, the model (2.1) generates a distinct type of bursting that depends on the activation of *I*_CAN_ through Ca^2+^ oscillations. We refer to this bursting as *(N)C-bursting*. The parentheses around ‘N’ indicate that the dependence of this bursting on *I*_NaP_ varies depending on *g*_NaP_ and *g*_Ca_. Specifically, for *g*_Ca_ ≥ 0.0004, and either 0 ≤ *g*_NaP_ ≤ 1.8 or *g*_NaP_ ≥ 3.5, the bursts are driven by *I*_CAN_ in the sense that they persist when *I*_NaP_ is blocked but disappear when *I*_CAN_ is blocked. This bursting type, known as *C-bursting*, represents a regime where the rhythmic bursting activity is maintained exclusively by *I*_CAN_. Within (N)C-bursting regime, there is also a subclass of bursting that depends on both *I*_NaP_ and *I*_CAN_, which we refer to as *NC-bursting*. This bursting exhibits two distinct subtypes: (Type 1) Both NaP and CAN mechanisms are essential for the NC bursting – blocking either channel abolishes the bursts. These bursts occur for 0.0001 ≤ *g*_Ca_ ≤ 0.00035 and 0 ≤ *g*_NaP_ ≤ 1.8 or *g*_NaP_ ≥ 3.5 within the red and orange regions in Figure 1(i); (Type 2) Bursting involves both *I*_NaP_ and *I*_CAN_, but the bursting continues if only one of the two currents is blocked. Bursting stops only when both channels are blocked. This type of (N)C-bursting occurs for *g*_Ca_ ≥ 0.0004 and 2 ≤ *g*_NaP_ ≤ 3 in Figure 1(i). In this paper, we focus on exploring the effects of NE on N- and C-bursters, excluding NC-bursters due to the limited experimental data on NE’s impact on those neurons, as a result of practical difficulties in isolating such pacemakers in experiments. For simplicity, we do not differentiate between C-bursting and the subtypes of NC-bursting in Figure 1(i), but instead distinguish (N)C-bursting types based on the timing of the elevation in cytoplasmic Ca^2+^ (i.e., *ca* jump-up) relative to burst initiation: the orange region corresponds to cases where bursting begins before the *ca* jump-up (Figure 1E), whereas the red region represents cases where *ca* jumps up before or near the onset of the burst (Figure 1F).

Finally, for intermediate values of *g*_Ca_, we observe *mixed bursting* (MB) patterns in the light orange region of Figure 1(i) (see Figure 1D for a representative trace). Such MB patterns, characterized by long bursts separated by sequences of short bursts, have been previously observed in preBötC models (e.g., [38, 31, 57, 58]), as well as in *in vitro* experiments, where the long burst can be associated with sighing [20, 35, 54, 40]. It remains unclear whether the MB solution shown in Figure 1D shares similar underlying mechanisms as those described in previous studies [57, 58]. Moreover, in Appendix D, we demonstrate that model (2.1) can generate a diverse range of MB patterns, some of which differ qualitatively from previously reported MB solutions and may exhibit quasi-periodic or chaotic dynamics (see Figure 13).

### 2.3. Modeling effects of norepinephrine (NE)

Neuromodulation is a process by which various substances influence neuronal activity. Many experimental and computational studies have made directed efforts into understanding the principles of neuromodulation, as well as how neuromodulators regulate respiratory rhythmic activity [19, 18, 52, 60, 56, 2, 48, 45, 26]. Experimental evidence has shown that neuromodulators like acetylcholine, serotonin, norepinephrine (NE), histamine, dopamine, and substance P can affect the frequency, amplitude, and regularity of respiratory activity (see [19] for review).

In mice, noradrenergic effects on respiration are considered to be primarily excitatory and mediated by *α*_1_-receptors [55]. In [56], NE was applied to synaptically isolated preBötC neurons in mice medullary brain slices, and the authors similarly found that the observed effects were mainly mediated by *α*_1_-receptor activation, despite NE also acting on other noradrenergic receptor subtypes. Further, the effects of NE on intrinsic bursting neurons (i.e., “pacemakers”) critically depended on whether their underlying bursting mechanisms were cadmium-insensitive (i.e., not dependent on calcium currents) or cadmium-sensitive (i.e., reliant on calcium currents). Specifically, NE increased the burst frequency, but did not change the burst duration in cadmium-insensitive bursters (e.g., N-bursting neurons). In contrast, in cadmium-sensitive bursters (e.g., (N)C-bursting neurons), the burst frequency did not change with NE, but the burst duration increased. Nonpacemakers also exhibited differential effects under the application of NE. In an “active nonpacemaker”, which continues to spike tonically when isolated from the network (e.g., a tonic spiking neuron), NE induced bursting. This induced pacemaking behavior was further found to be *I*_CAN_-dependent, as the NE-induced bursting activity continued in the presence of NaP-blocker riluzole, but was lost when *I*_CAN_ was blocked. In contrast, a “silent nonpacemaker”, which does not produce spikes when isolated from the network, continued to remain silent in the presence of NE.

[51] and [38] both modeled the effects of NE on the preBötC neurons based on that *α*_1_-receptor activation increases the CAN-current conductance in different cell types ([28], [27]). These models effectively captured several NE-mediated effects on preBötC neurons, such as the increase in burst frequency with unchanged burst duration in N-bursters and the increase in burst duration with unchanged burst frequency in C-bursters. However, none of these models account for the effects of NE on non-bursting neurons. A natural question arises: Can an increase in *g*_CAN_ also produce NE-induced effects on silent and tonic spiking neurons? The answer is no. We demonstrate in section 3 that increasing *g*_CAN_ alone does not induce a transition from spiking to bursting in single preBötC neuron models [51, 38, 31, 43]. In fact, bifurcation analysis in [57] suggests that increasing *g*_CAN_ can shrink the bursting region, favoring tonic spiking neurons - a result that is contrary to what has been observed in the experiments in [56]. Moreover, our computational simulations indicate that with a sufficient increase in *g*_CAN_, all silent neurons eventually transition to either a bursting or spiking behavior. Thus, while increasing *g*_CAN_ captures certain aspects of NE effects on bursting preBötC neurons, it does not fully capture all experimentally observed effects, particularly in non-bursting neurons.

In the review by [21], experiments in hepatocytes showed that IP_3_ functions as a second messenger for *α*_1_-adrenergic agonists and other calcium-mediated agonists. This was further supported by experiments in neurons [13]. Building on this, in addition to the previously described increases in *g*_CAN_, we propose that NE application in our model also leads to an increase in [IP_3_]. Incorporating this mechanism, we observe an overall increase in burst frequency in N-bursters as *g*_CAN_ and [IP_3_] increase (see Figure 3 in the Results section), consistent with the experimental findings in [56]. This increase is primarily driven by the enhanced depolarizing potential due to *g*_CAN_, while [IP_3_] has minimal impact on N-bursting dynamics. In C-bursters, burst frequency also increases with *g*_CAN_ (Figure 8) and we show in subsection 3.2 that this is mainly due to the involvement of the voltage-gated calcium current *I*_Ca_. However, we also demonstrate in subsection 3.2 that an increase in [IP_3_] may counteract the *g*_CAN_-induced frequency increase in the model (2.1), resulting in a constant burst frequency in the presence of NE, consistent with experimental observations. Nonetheless, there exist parameter regimes where increasing [IP_3_] and *g*_CAN_ leads to increases in both burst frequency and duration, which contradicts experimental results [56].

Different from previous NE modeling studies that overlook silent and tonic spiking neurons, a key novelty of our proposed mechanism is its ability to successfully capture NE-induced conditional pacemaking in tonic spiking neurons while also preserving the inactivity of silent neurons in the presence of NE. We show in subsection 3.3 that a tonic-spiking neuron can transition to bursting with increases in both *g*_CAN_ and [IP_3_]. Moreover, both mechanisms are essential: increasing *g*_CAN_ alone results in continued tonic spiking; while increasing [IP_3_] alone can induce bursting, these bursts are not C-bursts since they are lost when either *I*_NaP_ or *I*_CAN_ is blocked. Only bursts induced by simultaneous increases in *g*_CAN_ and [IP_3_] are truly CAN-dependent, as they persist when *I*_NaP_ is blocked but disappear when *I*_CAN_ is blocked - consistent with experimental data described above. This further confirms the necessity of incorporating both *g*_CAN_ and [IP_3_] increases to accurately model NE effects on preBötC neurons. Finally, we also identify a subset of silent neurons in our model that remains inactive despite increased *g*_CAN_ and [IP_3_] (see subsection 3.4), aligning with observations in [56].

In the following subsection, we provide a brief overview of geometric singular perturbation theory (GSPT) and the derivation of relevant subsystems and important geometric structures, which will be used in section 3 to analyze the distinct effects of NE on the different solution patterns of (2.1) as described above.

### 2.4. Geometric Singular Perturbation Theory

In addition to our modeling efforts, we apply techniques from dynamical systems theory, including phase plane analysis, fast-slow decomposition, and bifurcation analysis, to uncover mechanisms underlying different effects of NE on various types of intrinsic bursting preBötC neurons as well as the conditional bursting activity induced by NE. In this section, we use geometric singular perturbation theory (GSPT) [22, 46] to identify the underlying geometric structures that organize the dynamics of (2.1), which will be used for our analysis in section 3 of the mechanisms underlying NE modulation in individual preBötC neurons. For readers less familiar with GSPT and bifurcation analysis, we include a basic introduction to these mathematical approaches in Appendix A.

GSPT has been widely applied to study preBötC neurons, approached as either a two-timescale or a three-timescale problem [38, 57, 58, 59]. Traditionally, three-timescale problems were often simplified to two-timescale problems, which aligns with the natural setting of the GSPT approach [38, 3]. However, a two-timescale decomposition can fail to capture crucial dynamical aspects of a three-timescale system and is hence an insufficient approach for modeling and analysis of such problems [36, 41]. Our approach is to analyze model (2.1) as a three-timescale problem, based on our dimensional analysis detailed in Appendix C. This analysis transforms (2.1) to the following dimensionless system of equations:

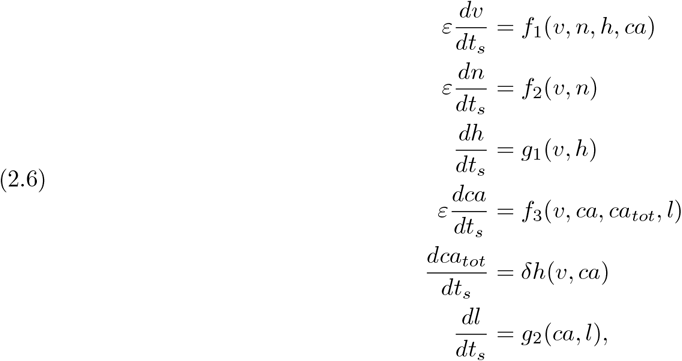

where *ε, δ* ≪ 1 are independent timescale parameters, *t*_*s*_ is the slow dimensionless time variable, *f*_1_, *f*_2_, *f*_3_, *g*_1_, *g*_2_, *h* are *O*(1) functions specified in Appendix B. The lowercase variables *v, ca, ca*_*tot*_ represent the dimensionless forms of *V*, [Ca] and [Ca]_Tot_, respectively, while *n, h* and *l* are already dimensionless. Throughout the paper, all model variables are plotted in their dimensionless forms. Time traces, however, are shown using the original time unit (ms).

From our analysis, we conclude that *v, n* and *ca* evolve on a fast timescale of *O*(*ε*^−1^), *h* and *l* evolve on a slow timescale of *O*(1), while *ca*_*tot*_ evolves on a superslow timescale of *O*(*δ*). We call system (2.6) that evolves over the *slow timescale t*_*s*_ the *slow system*, which is the same as (B.6) in Appendix C.

The existence of two independent singular perturbation parameters, *ε* and *δ*, implies there are various ways to implement GSPT, each yielding distinct singular limit predictions [36, 41], as detailed in Appendix C. In brief, applying GSPT yields the *fast layer problem* (see (C.3) in Appendix C), whose set of equilibrium points defines the *critical manifold M*_*s*_:

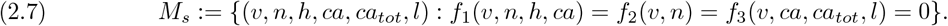

The slow evolution of trajectories of the *slow reduced problem* (see (C.6) in Appendix C) is slaved to *M*_*s*_ until nonhyperbolic points - such as the fold points *L*_*s*_ (defined in (C.5)) - are encountered.

Moreover, GSPT analysis also yields the *slow layer problem* (see (C.7) in Appendix C), whose equilibrium points form a one-dimensional subset of *M*_*s*_ called the *superslow manifold M*_*ss*_:

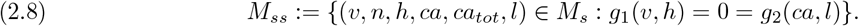

Near the singular limit *δ* → 0, the full solution trajectories travel near *M*_*ss*_ on the superslow timescale until they reach the nonhyperbolic points on *M*_*ss*_ - such as Hopf or saddle-node bifurcations (denoted as *L*_*ss*_) of the slow layer problem (C.7) - where Fenichel’s theory (GSPT) breaks down. For further details of these subsystems and their derivations, please refer to Appendix C. For brevity, we do not show the detailed constructions of singular orbits, which are solution segments of singular limit systems. Instead, we present orbits of the full system (2.6) and refer to different segments as being governed by various subsystems derived from the GSPT analysis, evolving under the fast, slow, or superslow flow. For further details on constructing singular orbit and using singular limit systems to understand the nature of the oscillations in the full system, see, e.g., [15, 41].

## 3. Results and Bifurcation Analysis

In this section, we use GSPT and bifurcation analysis to examine the dynamic mechanisms underlying the various activity patterns of model (2.6) as summarized in Figure 1(i) and analyze how changes in *g*_CAN_ and [IP_3_] replicate NE effects on preBötC neurons, as discussed in subsection 2.3. Specifically, we analyze the effects of NE on three distinct activity patterns of (2.6), using representative parameter values for each case: the *I*_NaP_-dependent N-bursting pattern with *g*_NaP_ = 2, *g*_CAN_ = 0.7, *g*_Ca_ = 0.00002 and [IP_3_] = 0.5 (subsection 3.1); the C-bursting pattern with *g*_NaP_ = 0, *g*_CAN_ = 0.7, *g*_Ca_ = 0.0005 and [IP_3_] = 0.5 (subsection 3.2); and the tonic spiking pattern with *g*_NaP_ = 4, *g*_CAN_ = 0.7, *g*_Ca_ = 0.0002 and [IP_3_] = 0.1 (subsection 3.3). We also identify a subset of silent preBötC neurons that remain inactive despite increased *g*_CAN_ and [IP_3_] in subsection 3.4. In the following subsections, we fix *g*_NaP_ and *g*_Ca_ for each solution pattern while varying *g*_CAN_ and [IP_3_] to investigate NE effects. All other parameters remain fixed as listed in Table 1.

### 3.1. Effects of NE on N-bursting neurons

The *I*_NaP_-dependent N-bursting dynamics can be generated by the voltage subsystem (*v, n, h*) without any calcium dynamics by setting *g*_CAN_ = 0. This type of bursting has been extensively studied using the fast-slow decomposition approach [46], by treating (*v, n*) as fast variables and *h* as a slow variable [38, 57, 59] (see our dimensional analysis in subsection 2.4). In the parameter regime for N-bursting in model (2.1), both *g*_Ca_ and [IP_3_] are low (see Figure 1(i)), keeping the calcium concentration *ca* at a low level. Although voltage dynamics influence the calcium compartment through *I*_Ca_, leading to small oscillations in *ca* around a low baseline, this effect remains negligible due to the low calcium levels. Similarly, the impact of [IP_3_] on the full dynamics via its effect on *ca* can also be neglected. As a result, *I*_CAN_, which is partially activated by *ca*, is functioning as a depolarizing leak current for *v*. In the following, we briefly review the bifurcation analysis for the N-burst and analyze the effects of NE on N-bursting dynamics by only considering the effect of increasing *g*_CAN_ on the voltage subsystem by examining its effect on the bifurcation diagram (see Figure 2).

**Fig. 2:**
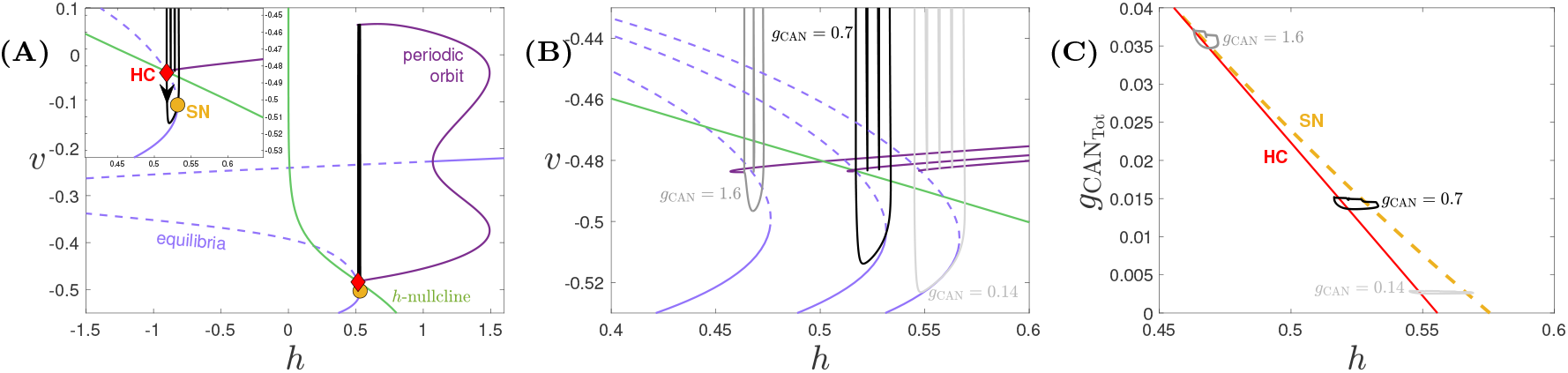
Projection of the *I*_NaP_-dependent N-burst solution of (2.6) along with the bifurcation diagrams for *g*_NaP_ = 2, *g*_Ca_ = 0.00002, [IP_3_] = 0.5, and varying values of *g*_CAN_. (A) *g*_CAN_ = 0.7. Projection of the solution (black) and the bifurcation diagram for the fast (*v, n*) subsystem with respect to *h*, along with the *h*-nullcline (green). The S-shaped light purple curve (solid where attracting, dashed otherwise) denotes the equilibria of the *v, n* equations and represents the projection of the critical manifold *M*_*s*_. The dark purple curves show the maximum and minimum *v* along two families of periodic orbits born at the subcritical Andronov-Hopf bifurcation (HB). The yellow circle and red diamond respectively denote the lower saddle-node (SN) bifurcation of *M*_*s*_ and the homoclinic (HC) bifurcation in which the outer periodic orbit branch terminates. (B): The effect of *g*_CAN_ on the solution trajectory and bifurcation diagram. From right to left, *g*_CAN_ = 0.14, 0.7, 1.6. (C): Projection of bursting solutions from panel (B) onto the 2-parameter bifurcation diagram in the (*h*,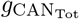)-space, where 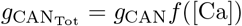 is given in (2.4).

Figure 2A shows the one-parameter bifurcation of the (*v, n*) subsystem with respect to *h*, together with the projection of the N-bursting solution (black trajectory) when *g*_NaP_ = 2, *g*_CAN_ = 0.7, *g*_Ca_ = 0.00002, [IP_3_] = 0.5 onto the (*h, v*)-space. The bifurcation diagram consists of an S-shaped curve of equilibria (S) and a family of stable periodic orbits (PO). The PO branch is born at a subcritical Andronov-Hopf bifurcation (HB) and terminates at a homoclinic bifurcation (HC). The equilibria curve S, which represents the projection of the critical manifold *M*_*s*_ onto the (*v, n*)-space, folds at two saddle-node (SN) bifurcations or fold points *L*_*s*_. The N-bursting solution is a square-wave burst that consists of a silent phase (i.e., interburst interval) along the lower branch of S and an active spiking phase along PO, initiating at SN and terminating at HC (Figure 2A inset).

Figure 2B shows the effect of increasing *g*_CAN_ on the bifurcation diagram from Figure 2A. As *g*_CAN_ increases, S and PO shift toward lower *h*, while the *h*-nullcline remains unchanged. The corresponding bursting trajectories at various *g*_CAN_ values are also superimposed onto (*h, v*)-space. Figure 2C summarizes how the key bifurcation points SN (yellow curve) and HC (red curve) move to lower *h* values with a continuous increase of *g*_CAN_ in the (*h*, 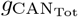)-space, where 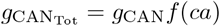 is a monotonically increasing function of *ca* (see (2.4)).

We first analyze the effect of *g*_CAN_ on the burst frequency by examining its influence on the interburst interval - the time between the end of one burst and the start of the next - which primarily determines the burst period [42]. For an N-bursting solution, this time interval can be approximated as

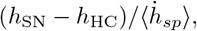

where 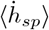 denotes the average speed *h* during the silent phase. As *g*_CAN_ increases, the lower fold of S shifts further away from the *h*-nullcline, leading to an increase in 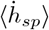. In the meantime, (*h*_SN_ − *h*_HC_) decreases with increasing *g*_CAN_, as the SN and HC curves move closer together and eventually merge in a saddle-node on an invariant circle (SNIC) bifurcation for *g*_CAN_ large enough. Together, the decrease in the numerator (*h*_SN_ − *h*_HC_) and the increase in the denominator 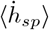 shorten the interburst interval, ultimately resulting in a higher burst frequency as *g*_CAN_ increases. This trend is confirmed by our numerical simulations in Figure 3(i), where *g*_CAN_ = 0.14, 0.7 and 1.6 correspond to the dark blue, light blue and orange regions. See also Figure 3(A), (B) and (D) for representative voltage traces, despite different [IP_3_] values.

**Fig. 3:**
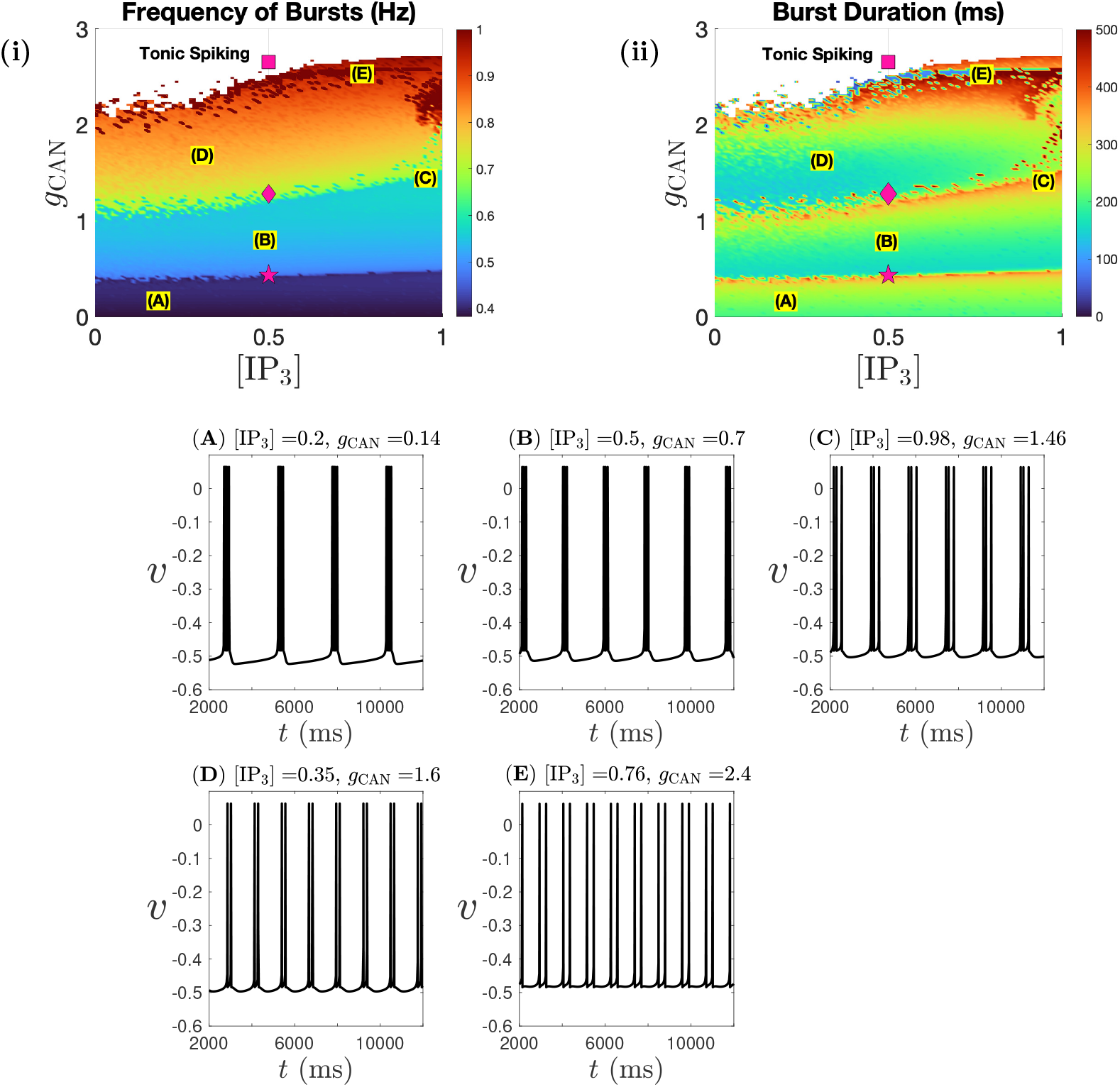
Two-parameter diagrams showing the effect of ([IP_3_], *g*_CAN_) on (i) N-burst frequency and (ii) N-burst duration for *g*_NaP_ = 2 and *g*_Ca_ = 0.00002. Pink symbols correspond to the *g*_CAN_ values at the saddle-node (SN) bifurcation points of the bursting branches in Figure 4 for fixed [IP_3_] = 0.5. Sample voltage traces corresponding to labeled parameter values (A) through (E) are displayed below.

Notably, the burst duration remains similar across these three regions. To understand this, we switch to examining the active spiking phase within the burst. During this phase, the trajectory oscillates between the maximum and minimum *v* branches along the family of periodic orbits (PO). As *g*_CAN_ increases, the lower *v* branch shifts to lower *h* values while the *h*-nullcline remains static (see Figure 2B). As a result, the trajectory transitions from oscillating entirely above the *h*-nullcline (e.g., *g*_CAN_ = 0.14) to a state where part of the trajectory during the active phase falls below the *h*-nullcline (e.g., *g*_CAN_ = 1.6). In other words, as *g*_CAN_ increases, a larger portion of the full system trajectory’s projection in the (*h, v*)-space during the active phase lies close to or below the *h*-nullcline. Consequently, the average rate of change of *h* during the active phase becomes less negative (i.e., the spiking frequency within the burst decreases). Meanwhile, as discussed above, the distance (*h*_SN_ − *h*_HC_) decreases with increasing *g*_CAN_. These two effects counterbalance each other, maintaining a nearly constant burst duration, as confirmed by our numerical simulations in Figure 3(ii), where the burst duration remains constant in most green regions. However, in certain regions, where the reduced *h* distance can no longer fully offset the slower *h* rate, the burst duration increases as indicated by the red regions (also compare Figure 3(B) and (C)).

The effect of [IP_3_] on calcium dynamics has been analyzed previously in [57] in the absence of *I*_Ca_. Specifically, increasing [IP_3_] triggers greater release of stored Ca^2+^ from the ER into the cytoplasm, thereby raising intracellular calcium levels. This phenomenon largely holds in our model, despite the presence of *I*_Ca_ which remains small in the NB parameter regime. During the active phase, Ca^2+^ enters the cell through *I*_Ca_, contributing to neuronal depolarization via *I*_CAN_. However, during the interburst interval, calcium quickly returns back to its low baseline level by calcium pumps and thus has minimal impact on voltage dynamics compared to the effect of *g*_CAN_ (see Figure 3).

Furthermore, our analysis suggests that the increase in burst frequency is neither continuous nor strictly monotonic but instead occurs in a discrete fashion, exhibiting a phasic pattern. Specifically, Figure 3(i) shows that increasing *g*_CAN_ has minimal effects on burst frequency over a relatively broad parameter range, but can lead to a sudden, significant increase in burst frequency upon crossing a threshold. Similarly, discrete changes in burst duration are observed across the same threshold curves (Figure 3(ii)). To better understand these thresholds, we analyze bifurcations of the full system (2.6) with respect to *g*_CAN_ at fixed [IP_3_] = 0.5. Figure 4 shows the effect of *g*_CAN_ on the frequency of different bursting branches (dark blue, light blue and yellow curves — color-coded to match the burst frequency values in Figure 3(i)) and the tonic spiking branch (black). As *g*_CAN_ increases, the frequency of each N-burst branch remains nearly constant until its saddle-node (SN) bifurcation is reached. After crossing this point, the solution transitions to the next branch, resulting in a discrete jump in burst frequency. After crossing the SN of the yellow burst branch at the pink square, the solution jumps to the tonic spiking branch, leading to tonic spiking solutions for *g*_CAN_ large enough. These three SN bifurcation points are overlaid on the ([IP_3_], *g*_CAN_)-space in Figure 3 and align with the three threshold curves, suggesting that the discrete changes in the N-burst frequency are driven by the saddle-node bifurcations of the N-burst solution branches. Another interesting observation in Figure 4 is the loss of a spike within each burst as the solution crosses each SN bifurcation threshold with increasing *g*_CAN_ (see also Figure 2B). This is because, as *g*_CAN_ increases, the burst duration remains almost constant while the intra-burst spike frequency decreases. As a result, the system can no longer sustain the original number of spikes per burst, leading to a gradual loss of spikes as SN points are crossed.

**Fig. 4:**
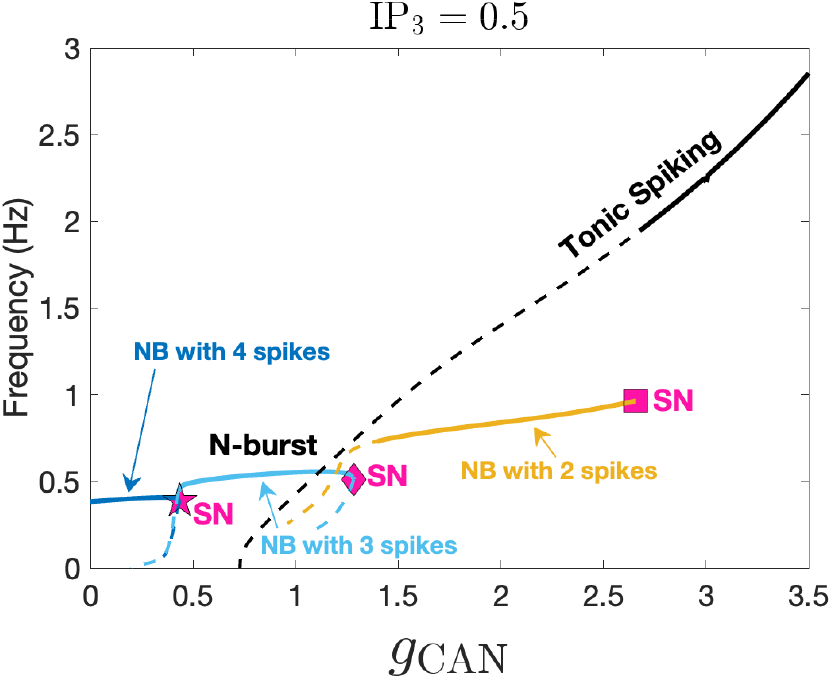
Bifurcation diagram of (2.6), showing the frequency of different N-burst solutions (colored curves) and the tonic spiking branch (black curve) as functions of *g*_CAN_. As *g*_CAN_ increases, the number of spikes per burst decreases from 4 (dark blue) to 3 (light blue), then to 2 (yellow), before eventually transitioning to tonic spiking. Solid lines denote stable branches; dashed lines denote unstable branches. All parameter values are as in Figure 2A.

To summarize, our analysis and simulations of (2.6) predict that increasing *g*_CAN_ increases the burst frequency of N-bursting solutions through a sequence of SN bifurcations, while having minimal impact on burst duration. Additionally, [IP_3_] has little effect on both burst frequency and duration. The agreement with experimental data described before in subsection 2.3 supports our proposed mechanism for NE, which involves increasing *g*_CAN_ and [IP_3_]. Although it seems that IP_3_ is not a prerequisite for modeling the effect of NE on N-bursts, we show later that it is critical to capture the effects of NE on other cell types of the preBötC.

### 3.2. Effect of NE on C-bursting neurons

When *g*_NaP_ = 0, *g*_CAN_ = 0.7, *g*_Ca_ = 0.0005 and [IP_3_] = 0.5, model (2.6) produces C-bursting dynamics (see Fig. Figure 1(i), lower right red region, label (F); also reproduced in Figure 5A). Below, we first apply GSPT analysis (see subsection 2.4) to briefly explain the underlying mechanism for the C-burst, by considering the projection of the critical manifold *M*_*s*_, superslow manifold *M*_*ss*_, and the solution trajectory onto the (*ca, ca*_*tot*_, *l*)-space (Figure 5B). Then we examine the effect of *g*_CAN_ and [IP_3_] on the C-bursting dynamics by considering their influence on the geometric structures of the calcium subsystem (see Figures 6 and 7).

**Fig. 5:**
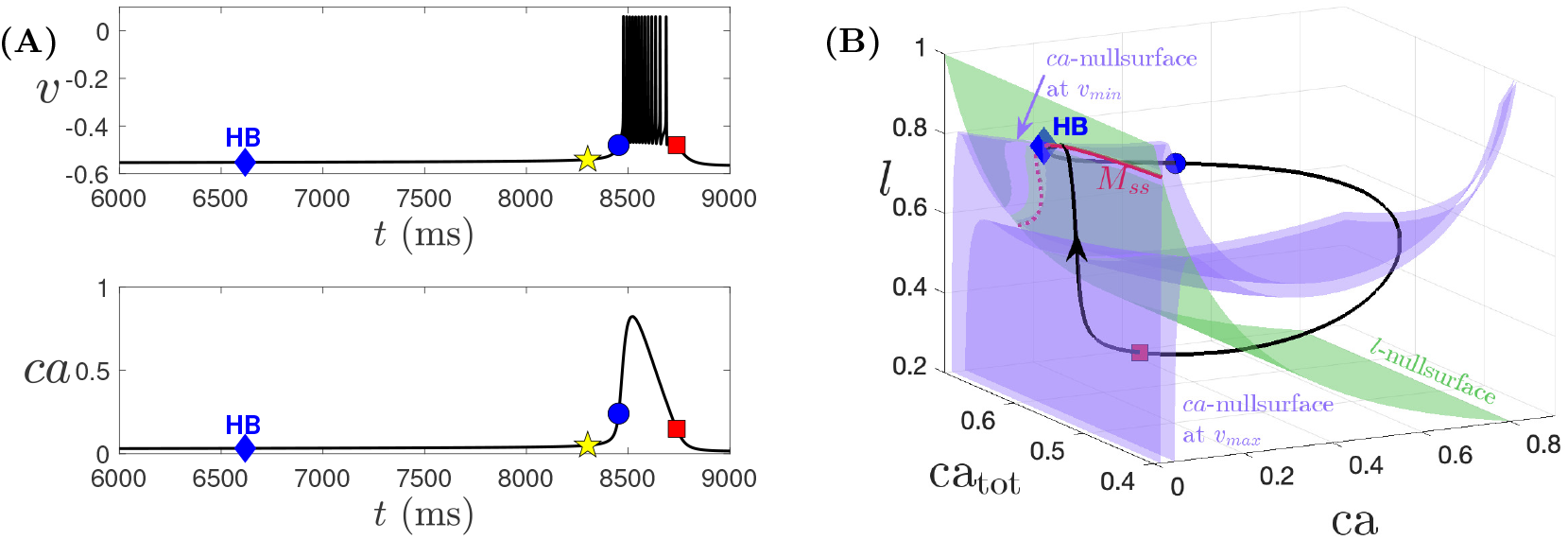
Simulation of one cycle of the C-bursting solution generated by (2.6) for *g*_NaP_ = 0, *g*_CAN_ = 0.7, *g*_Ca_ = 0.0005 and [IP_3_] = 0.5, and other parameters at default values. The blue diamond denotes the HB point. The yellow star denotes the jump up point of *ca*, defined as crossing the threshold of *ca* = 0.05 from below, whereas the blue circle and red square mark the initiation and termination of the burst. (A) Temporal evolution of *v* and *ca*. (B) Projections of *M*_*s*_ (i.e., *ca*-nullsurfaces denoted by blue surfaces) with *v* at its minimum and maximum in the (*ca, ca*_*tot*_, *l*)-space, together with the black solution trajectory from panel (A) and the superslow manifold *M*_*ss*_ (red curve). The green surface denotes the *l*-nullsurface.

**Fig. 6:**
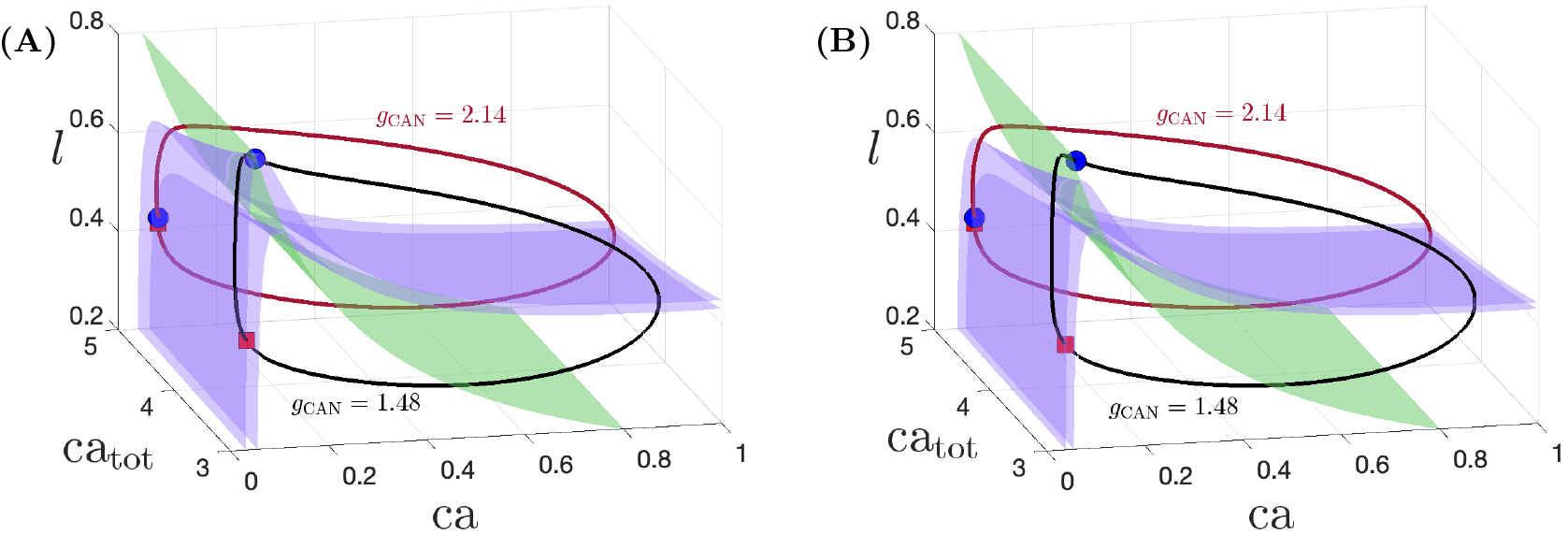
*ca*-nullsurfaces (blue surface) and *l*-nullsurface (green surface) for [IP_3_] = 0.2 and (A) *g*_CAN_ = 1.48 and (B) *g*_CAN_ = 2.14, projected onto (*ca, ca*_*tot*_, *l*)-space. Solution trajectories of (2.6) with *g*_CAN_ = 1.48 and *g*_CAN_ = 2.14 are shown in both panels by the black and red curves, respectively. Other parameters, color coding and symbols are the same as Figure 5B.

**Fig. 7:**
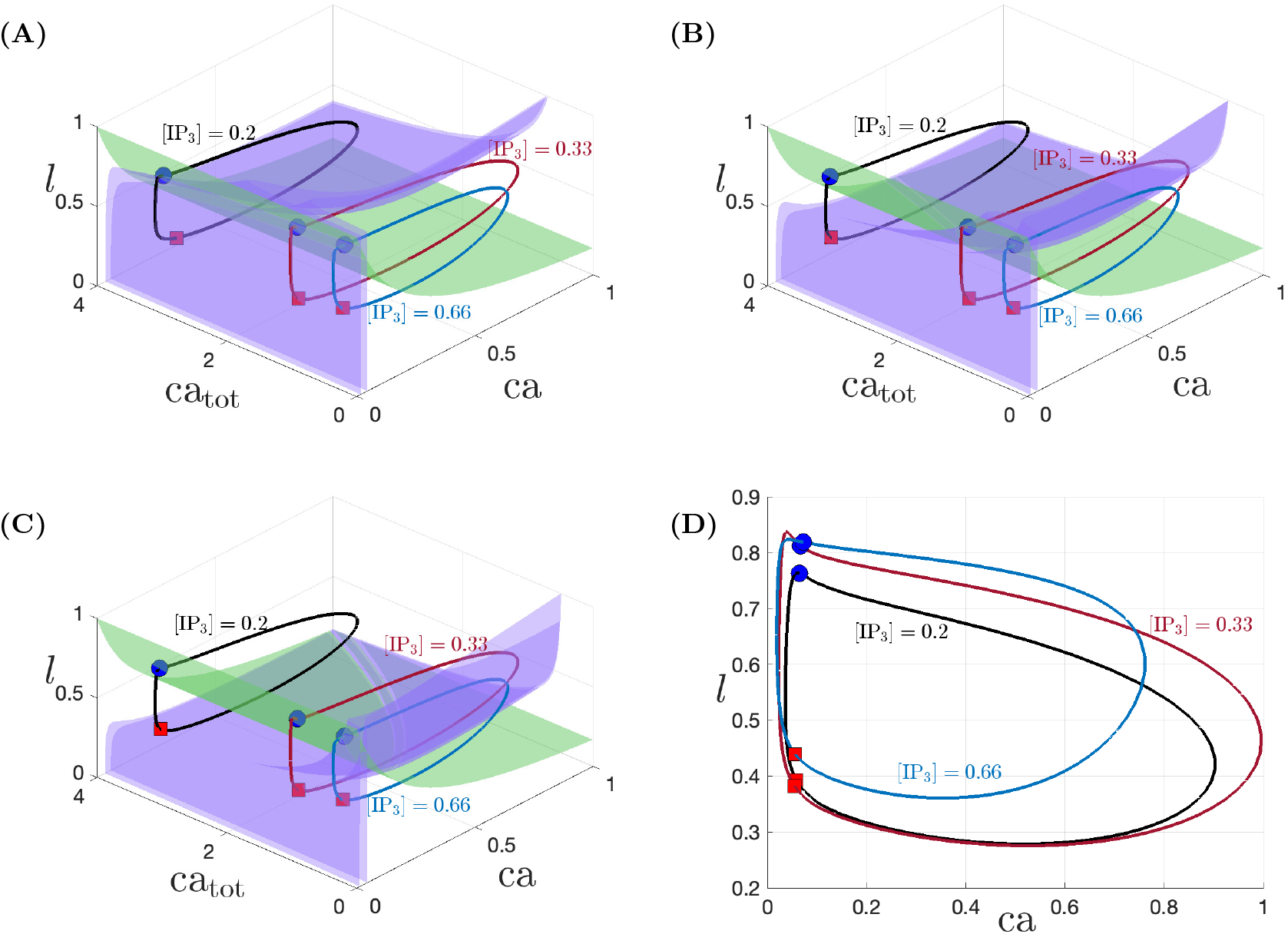
*ca*-nullsurfaces and *l*-nullsurface for *g*_CAN_ = 1.48 and (A) [IP_3_] = 0.2, (B) [IP_3_] = 0.33 and (C) [IP_3_] = 0.66, projected onto (*ca, ca*_*tot*_, *l*)-space, and (D) onto (*ca, l*)-space. Solution trajectories of (2.6) with [IP_3_] = 0.2, [IP_3_] = 0.33 and [IP_3_] = 0.66 are shown in all panels by the black, red and blue curves, respectively, in all panels. Other parameters, color coding and symbols are the same as in Figure 5B.

Past studies have shown that C-bursting is driven by *ca*-oscillations [47, 38, 57]. Indeed, Figure 5(B) illustrates that the *ca*-oscillation begins at the yellow star (defined as crossing *ca* = 0.05 from below), preceding the burst onset at the blue circle. To analyze this mechanism, we apply the GSPT approach, which starts with a bifurcation analysis of the fast layer problem (*v, n, ca*) while treating the slow variables (*h, ca*_*tot*_, *l*) as bifurcation parameters. This requires a visualization of the critical manifold *M*_*s*_, defined by *f*_1_(*v, n, h, ca*) = *f*_2_(*v, n*) = *f*_3_(*v, ca, ca*_*tot*_, *l*) = 0 (see (2.7)). Since *g*_NaP_ remains small in the C-bursting parameter regime, the bifurcation diagram for the fast voltage subsystem, discussed in the previous section, is no longer relevant. Instead, following [58], we examine the trajectory and *M*_*s*_ in (*ca, ca*_*tot*_, *l*)-space to understand the dynamics of calcium, which depends on the neuronal membrane potential *v* through *I*_Ca_. A projection onto (*v, n, ca*) can then be used to understand the C-bursting dynamics in the full system, which is driven by *ca*-oscillations via *I*_CAN_ activation. In this paper, we focus only on the former projection for analyzing the *ca*-oscillations and the effects of NE, referring interested readers to [58] for further details.

Since *f*_3_(*v, ca, ca*_*tot*_, *l*) does not depend on *h*, we can visualize the projection of *M*_*s*_ onto (*ca, ca*_*tot*_, *l*)-space by considering *ca*-nullsurfaces, each defined for *v* fixed (see Figure 5B, the two blue surfaces). Also shown are the projections of the solution trajectory (black), the superslow manifold *M*_*ss*_ (red), and the *l*-nullsurface (green). After the burst ends at the red square in Figure 5B, the trajectory jumps to the left branch of the *ca*-nullsurface at *v*_*min*_, entering the silent phase. It then evolves on the slow timescale under the slow reduced layer problem (C.10), with the superslow variable *ca*_*tot*_ remaining approximately constant, until reaching the stable portion of the superslow manifold *M*_*ss*_ (red curve). After that, the trajectory transitions to the superslow timescale under the superslow reduced problem (C.9). The trajectory closely follows the attracting side of *M*_*ss*_, passes over the HB (blue diamond) to the repelling side of *M*_*ss*_, and undergoes a delay in which it traces the repelling branch before making a fast jump to large *ca* (see Figure 5A, between blue diamond and yellow star). This jump depolarizes the membrane potential sufficiently through *I*_CAN_ activation, triggering burst onset in *v* in the C-bursting solution. After the calcium jump, the trajectory slowly travels along the right branch of *ca*-nullsurface before passing its lower fold and making a fast jump back to the red square, terminating the burst and completing a full cycle.

Next we investigate how increasing *g*_CAN_ and [IP_3_] affects the C-burster by examining the projections of solution trajectories and *M*_*s*_ onto (*ca, ca*_*tot*_, *l*) (see Figures 6 and 7). Additionally, we numerically simulated the effects of *g*_CAN_ and [IP_3_] on burst frequency and duration, as shown in Figure 8(i) and (ii). Under fixed [IP_3_] = 0.2, increasing *g*_CAN_ depolarizes the cell, which increases the total calcium concentration within the cell through *I*_Ca_. As a result, the solution for higher *g*_CAN_ lies at higher *ca*_*tot*_ values than a solution for low *g*_CAN_ (see Figure 6). The projections of *ca*-nullsurfaces for *g*_CAN_ = 1.48 and 2.14 are shown respectively in panels (A) and (B). From our analysis of C-bursting, the burst period is determined by the *ca*-oscillation period, which is largely set by the trajectory’s duration along the left branch of the *ca*-nullsurface at *v*_*min*_. In contrast, the burst duration is determined by the duration of the active phase when *ca* is relatively large (Figure 5, the right segment of the trajectory between blue circle and red square). Therefore, to examine the effect of NE on C-burst frequency and duration, we analyze different phases of the *ca*-oscillation.

**Fig. 8:**
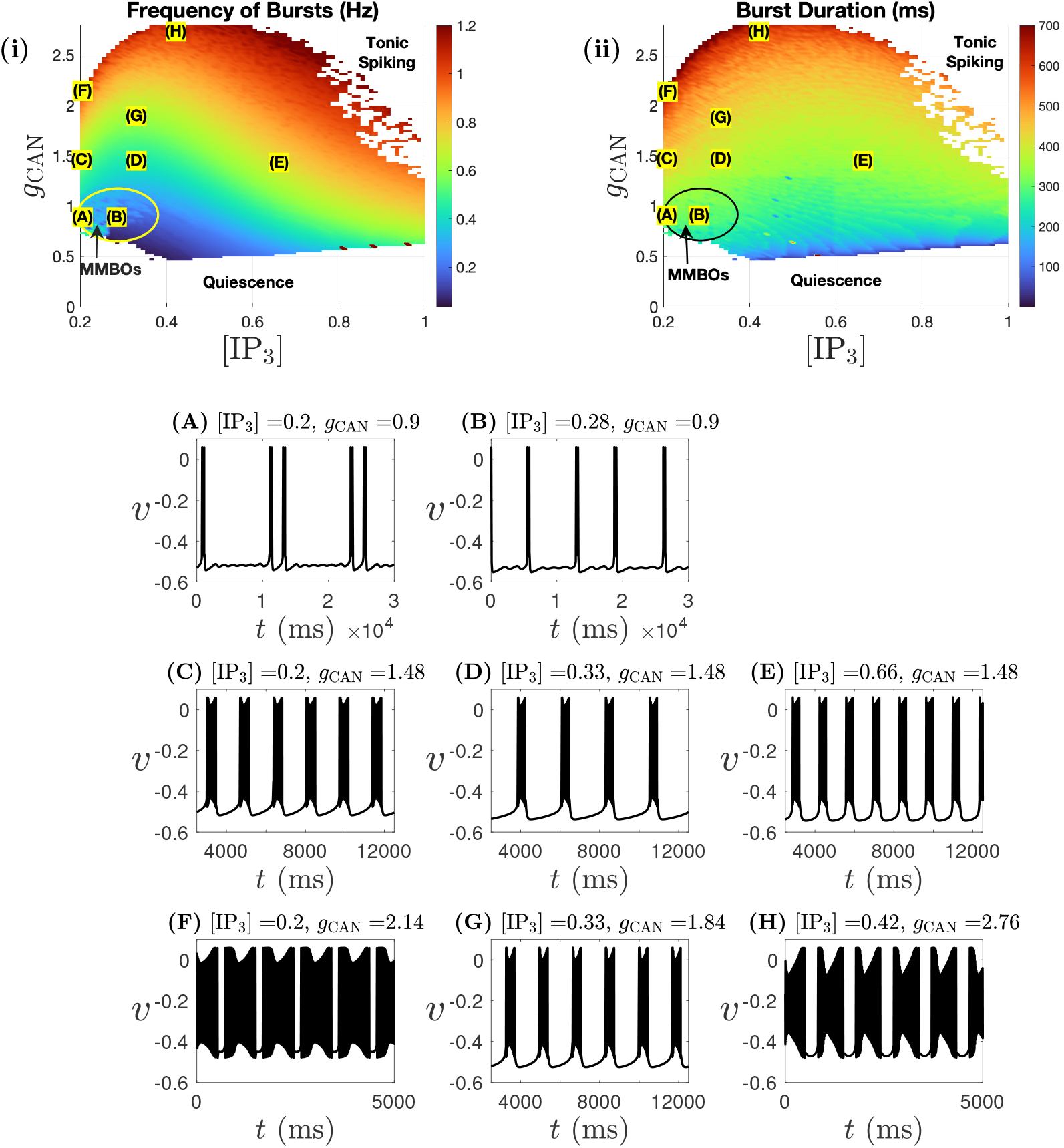
Two-parameter diagrams showing the effects of ([IP_3_], *g*_CAN_) on (i) C-burst frequency and (ii) C-burst duration. Sample voltage traces corresponding to labeled parameter values (A) through (H) are displayed below.

Beginning at the red square in Figure 6A, the trajectory for *g*_CAN_ = 1.48 (black curve) follows the left branch of the *ca*-nullsurface at *v*_*min*_ (upper blue surface) until it reaches the fold, then undergoes a fast jump to the right branch, initiating burst onset. Here, since the jump-up of *ca* occurs at HB which lies very close to the fold of the *M*_*s*_, we do not include the *M*_*ss*_ and the HB in the figure. As *g*_CAN_ increases, the trajectory (red curve) shifts to higher *ca*_*tot*_ and moves further away from the *l*-nullsurface. At the same time, the *ca*-nullsurface (blue surfaces in Figure 6B) shifts slightly toward lower *l*-values. As a result, the rate of increase of *l* during the silent phase becomes slightly higher and the red trajectory crosses the fold of the *ca*-nullsurface at a lower *l*-value. Together, these effects result in faster ca-oscillations and an increased C-burst frequency. On the other hand, a higher *g*_CAN_ lowers the *ca*-threshold required to trigger a *v*-spike in the voltage subsystem, advancing the onset of the C-burst and significantly shortening the silent phase duration. Meanwhile, the trajectory spends more time in the active phase of the cycle, allowing the neuron to burst for a longer duration. These analysis is consistent with our numerical simulations, which show that increasing *g*_CAN_ for fixed [IP_3_] consistently increases both burst frequency and duration (Figure 8).

Figure 7 shows the effect of increasing [IP_3_] on the C-bursting dynamics for fixed *g*_CAN_ = 1.48. The projections of the *ca*-nullsurfaces for [IP_3_] = 0.2, [IP_3_] = 0.33 and [IP_3_] = 0.66 are shown respectively in panels (A), (B) and (C). The solution trajectories for all three [IP_3_] values are shown in all panels. Comparing panels (A-C) indicates that increasing [IP_3_] shifts the *ca*-nullsurface towards lower *l*-values while pushing the trajectory to lower *ca*_*tot*_ values. This occurs because higher [IP_3_] enhances calcium release from the ER into the cytosol, which in turn increases calcium extrusion out of the cell through plasma membrane Ca^2+^ pumps, reducing the total intracellular calcium.

We first compare [IP_3_] = 0.2 and 0.33, with their trajectories also projected onto (*ca, l*)-space (Figure 7D). This projection clearly shows that the C-burst for [IP_3_] = 0.33 initiates at a higher *l* value than for [IP_3_] = 0.2, while both bursts terminate at similar *l* values (red square). To understand why, we examine the projection onto (*ca, ca*_*tot*_, *l*)-space in panels (A) and (B). For both [IP_3_] values, burst onset (blue circle) occurs approximately along the fold curve of the *ca*-nullsurface. While increasing [IP_3_] shifts the *ca*-nullsurface to lower *l* values, it also significantly lowers the trajectory’s *ca*_*tot*_, which in turn increases the *l* value at the *ca*-nullsurface’s fold. As a result, the jump up of *ca* for [IP_3_] = 0.33 occurs at a higher value of *l* than for [IP_3_] = 0.2 (see Figure 7D, blue circles). Similarly, if the *ca*-nullsurface remained fixed while the trajectory alone shifted to lower *ca*_*tot*_, one would expect a higher *l* value at burst termination (red square) as the right branch of the *ca*-nullsurface moves to higher *l* as *ca*_*tot*_ decreases. Nonetheless, since the increased [IP_3_] also shifts the surface to lower *l*, these two effects counteract each other, leading to similar *l* values at burst termination. Hence, as a result of the evolution rate of *l* remaining similar, the longer distance between the blue circle and red square along the left branch of *M*_*s*_ at [IP_3_] = 0.33 results in a longer interburst interval, thereby reducing burst frequency compared to [IP_3_] = 0.2 (see Figure 8C and D).

A similar argument also holds when comparing [IP_3_] = 0.33 and [IP_3_] = 0.66. While increasing [IP_3_] continues to push the left branch of *ca*-nullsurface to lower *l* (compare Figure 7B and C), the effect is less pronounced than in the previous comparison. As a result, the two opposing effects described above largely cancel out near the left branch, resulting in a similar *l* value at burst onset (blue circles). However, the jump-back value of *l* (red square) is higher for [IP_3_] = 0.66 than for [IP_3_] = 0.33, as the right branch is only minimally affected by the increased [IP_3_] but shifts to higher *l* with larger *ca*_*tot*_ along the blue trajectory. Consequently, the C-burst for [IP_3_] = 0.66 has a higher burst frequency than for [IP_3_] = 0.33 (see Figure 8D and E). This suggests a non-monotonic effect of [IP_3_] on the C-burst frequency, where an initial increase in [IP_3_] slows down burst frequency, but a further increase accelerates it (see Figure 8(i)). We omit a detailed analysis of the effect of [IP_3_] on burst duration, which requires a closer examination of the trajectory along the right branch, its shape variations, and the average rate of *ca* during the burst. It is worth noting that the effect of [IP_3_] on the size of the (*ca, l*)-projection of the trajectory during the burst is also non-monotonic (see Figure 7D).

Having analyzed the effects of *g*_CAN_ and [IP_3_] separately, we now examine their combined influence. Figure 8 suggests that increasing both *g*_CAN_ and [IP_3_] generally leads to an increase in the C-burst frequency. However, as previously analyzed, while increasing *g*_CAN_ consistently raises burst frequency, an increase of [IP_3_] from a relatively low level initially reduces it. Thus, in theory, there should exist a parameter region where these two opposing effects counterbalance, resulting in a constant burst frequency, as observed in experiments. Indeed, Figure 8C and G show that the C-burst frequency remains nearly unchanged despite a 20% increase in *g*_CAN_, due to the opposing effect of simultaneously increasing [IP_3_]. A similar example is seen between Figure 8F and H. Similarly, as a result of the opposing effects of *g*_CAN_ and [IP_3_] on the burst duration, as seen in Figure 8(ii), the combined influence leads to one of the two outcomes: (1) the increase in C-burst duration caused by *g*_CAN_ is counterbalanced by the decrease induced by [IP_3_], resulting in little change or a slight decrease in duration (see Figure 8C-G for example); or (2) C-burst duration increases (see Figure 8G-H, note the different time scale).

Finally, we note that model (2.6) exhibits *mixed-mode bursting oscillations* (MMBOs) in a small region of parameter space (Figure 8(i), yellow circle; see also, Figure 8A,B). These oscillations exhibit complex dynamics characterized by alternating large-amplitude bursts and small-amplitude oscillations [16, 17]. While not central to our present modeling study, the emergence of MMBOs highlights the richness of the model’s dynamics, and investigating their underlying mechanism represents an interesting direction for future work.

### 3.3. NE-induced Bursting in a Tonic Spiking neuron

We next model the CAN-dependent conditional bursting behavior evoked in tonic spiking neurons as experimentally observed by [56]. In model (2.6) with *g*_CAN_ = 0.7 as the control condition, tonic spiking arises under two distinct parameter regimes: either when calcium conductance (*g*_Ca_) is very small (less than 0.0001) (see Figure 1(i), light blue region), or when *g*_Ca_ is intermediate (e.g., ∼ 0.0002) (see Figure 9(i), light blue region). In both cases, *g*_NaP_ needs to be relatively high to sustain tonic spiking in the voltage dynamics.

**Fig. 9:**
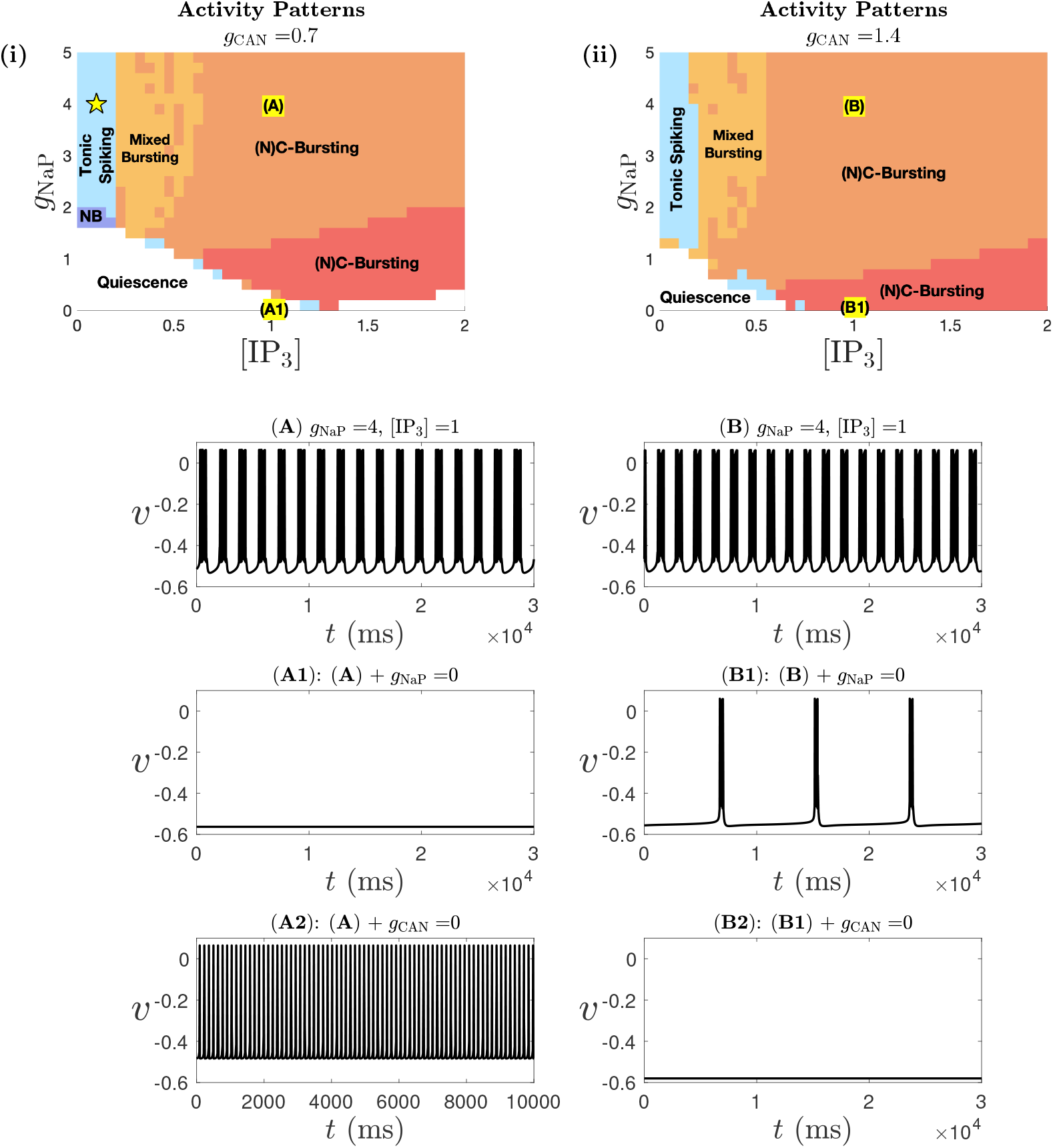
Activity patterns of model (2.6) for *g*_Ca_ = 0.0002, shown as a function of *g*_NaP_ and [IP_3_], with (i) *g*_CAN_ = 0.7 and (ii) *g*_CAN_ = 1.4. Color coding follows that in Figure 1(i). (A): A sample voltage trace demonstrates that increasing [IP_3_] induces rhythmic bursting activity corresponding to point (A) in panel (i); this bursting is abolished when either *I*_NaP_ (A1) or *I*_CAN_ (A2) is blocked. (B): A sample voltage trace for NE (*I*_CAN_+IP_3_)-induced bursting. This bursting persists in the absence of *I*_NaP_ (B1), but is eliminated when *I*_CAN_ is also blocked (B2).

We first focus on the tonic spiking regime highlighted in Figure 9(i), characterized by intermediate *g*_Ca_ and relatively low [IP_3_]. Our simulations and analyses highlight three important observations: First, neurons in this regime remain tonic spiking as *g*_CAN_ increases from 0.7 to 1.4 (compare Figure 9(i) and (ii)) and even higher values (data not shown), suggesting that increasing *g*_CAN_ alone is insufficient to reproduce NE-induced bursting observed experimentally. Second, while increasing [IP_3_] alone does induce bursting (Figure 9A), this activity is not solely dependent on CAN currents. Third, when both *g*_CAN_ and [IP_3_] are increased, the system transitions from tonic spiking to bursting (Figure 9B), with the bursting behavior solely dependent on CAN currents, consistent with experimental data. These observations suggest that, within this parameter regime, coordinated modulation of both *g*_CAN_ and [IP_3_] is essential to reproduce the experimentally observed NE-induced, CAN-dependent-only bursting from tonic spiking neurons.

To illustrate this, we show voltage traces from a representative control tonic spiking neuron with *g*_NaP_ = 4, *g*_Ca_ = 0.0002, [IP_3_] = 0.1 and *g*_CAN_ = 0.7 (yellow star in Figure 9(i)) after it transitions to bursting, either through an increase in [IP_3_] alone or under combined increases in both *g*_CAN_ and [IP_3_]. Importantly, the IP_3_-induced bursting corresponding to point (A) in Figure 9(i), with its voltage trace shown in Figure 9A, depends on both *I*_NaP_ and *I*_CAN_. Blocking either current eliminates bursting (see Figure 9A1 and A2), contradicting experimental evidence suggesting that NE-induced bursting is purely CAN-dependent. In contrast, bursting induced by simultaneously increasing *g*_CAN_ and [IP_3_], corresponding to point (B) in Figure 9(ii), with its voltage trace shown in Figure 9B, is a C-burst. This bursting persists in the absence of *I*_NaP_ (Figure 9B1) but is abolished when *I*_CAN_ is blocked (Figure 9B2). Below we refer to the bursting in Figure 9A as IP_3_-induced bursting, and the bursting in Figure 9B as NE-induced bursting.

Our GSPT analysis of the NE-induced bursting (Figure 10) reveals a bursting mechanism distinct from that of the regular C-burst in subsection 3.2. The corresponding temporal evolution of *v* and *ca* over one full burst cycle is shown in panel A. The blue circle and red square mark the initiation and termination of the burst, respectively, corresponding to the points where the trajectory projected onto (*ca, h*)- and (*ca, h, v*)-spaces crosses the fold *L*_*s*_ of the critical manifold *M*_*s*_ (see Figure 10B and its inset). During the silent phase, the trajectory follows the attracting lower branch of *M*_*s*_ until it reaches its fold *L*_*s*_ at the blue circle, triggering burst initiation and entering the active phase. This in turn causes *ca* to jump up. As *h* and *ca* decrease, the trajectory again crosses *L*_*s*_ - now at the red square - which coincides with a homoclinic bifurcation at a saddle-node on invariant circle (SNIC) bifurcation that terminates the periodic branch of the fast layer problem. Thus, the burst ends at the red square. This bursting mechanism clearly differs from the C-burst described in subsection 3.2, where bursts are initiated as *ca* jumps up near a HB along *M*_*ss*_. Nonetheless, the fact that *ca*-oscillations do not initiate the NE-induced burst does not imply they are not important. The jump-up of *ca* helps pull the trajectory away from the fold *L*_*s*_ after crossing the blue circle, leading to continuous spiking during the active phase. Moreover, the subsequent decrease of *ca* is crucial for burst termination - without it, the system would remain in a tonic spiking state.

**Fig. 10:**
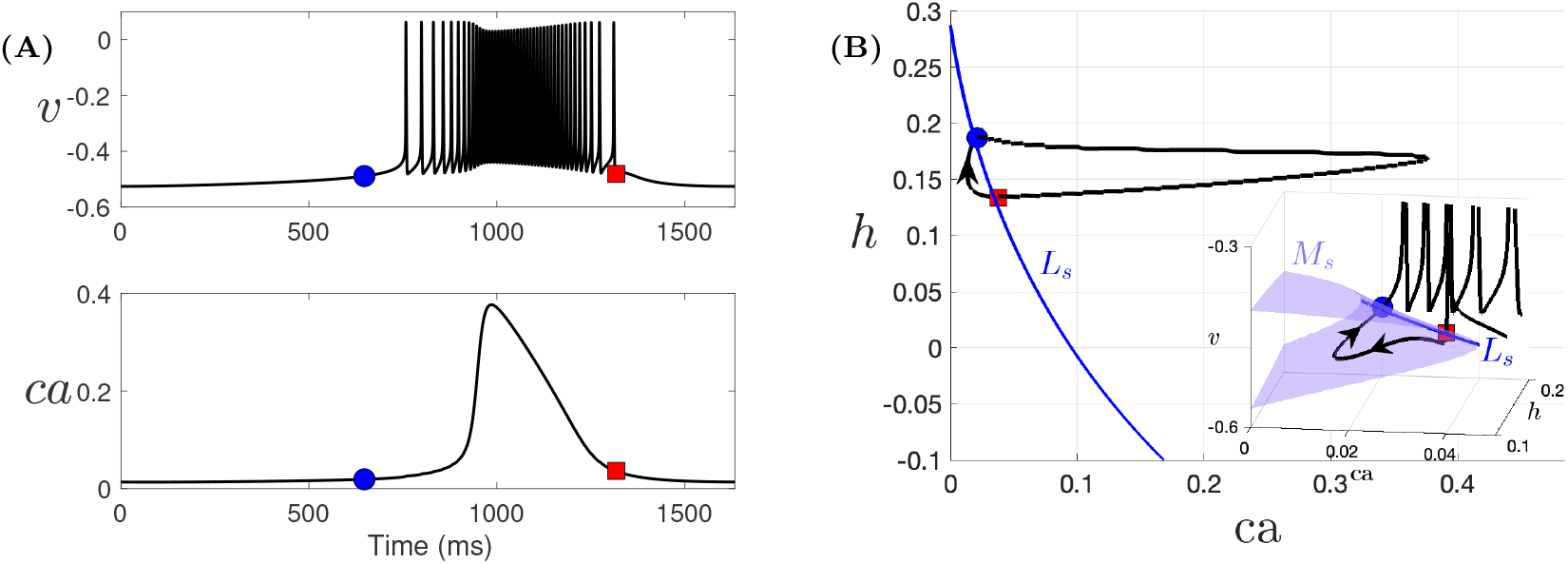
Simulation of one cycle of NE-induced C-bursting solution from Figure 9B. The blue circle and red square mark the initiation and termination of the burst. (A) Temporal evolution of *v* and *ca* over one full cycle. (B) Projection of the black trajectory from panel (A) onto the (*ca, h*)-space, along with the fold *L*_*s*_ (blue curve) of the critical manifold *M*_*s*_. The inset shows the trajectory, *M*_*s*_ (blue surface), and *L*_*s*_ projected onto (*ca, h, v*)-space. The burst is initiated and terminated as the trajectory crosses *L*_*s*_.

To better understand why blocking *g*_NaP_ eliminates IP_3_-induced bursting while NE-induced bursting persists, we examine the projections of the dynamics with *g*_NaP_ = 0 from Figure 9A1 and B1 onto (*ca, ca*_*tot*_, *l*)-space, together with the superslow manifold *M*_*ss*_ (see Figure 11A and B). In panel (A), the system settles into a stable equilibrium, corresponding to a silent state. This explains why IP_3_-induced bursting becomes silent in the absence of *I*_NaP_. In contrast, in panel (B) with a higher *g*_CAN_ value, the equilibrium is unstable and a bursting solution emerges. The bursting trajectory initially travels along the stable branch of the *ca*-nullsurface, then transitions onto the stable portion of *M*_*ss*_, traveling along it until it reaches the fold *L*_*ss*_, after which it jumps to higher *ca* values. This *ca* jump triggers burst onset at the blue circle. The mechanism is similar to that of the regular C-burst described in subsection 3.2 as both are driven by calcium oscillations; however, one is initiated by crossing the fold *L*_*ss*_, while the other is triggered by a Hopf bifurcation and experiences a delay.

**Fig. 11:**
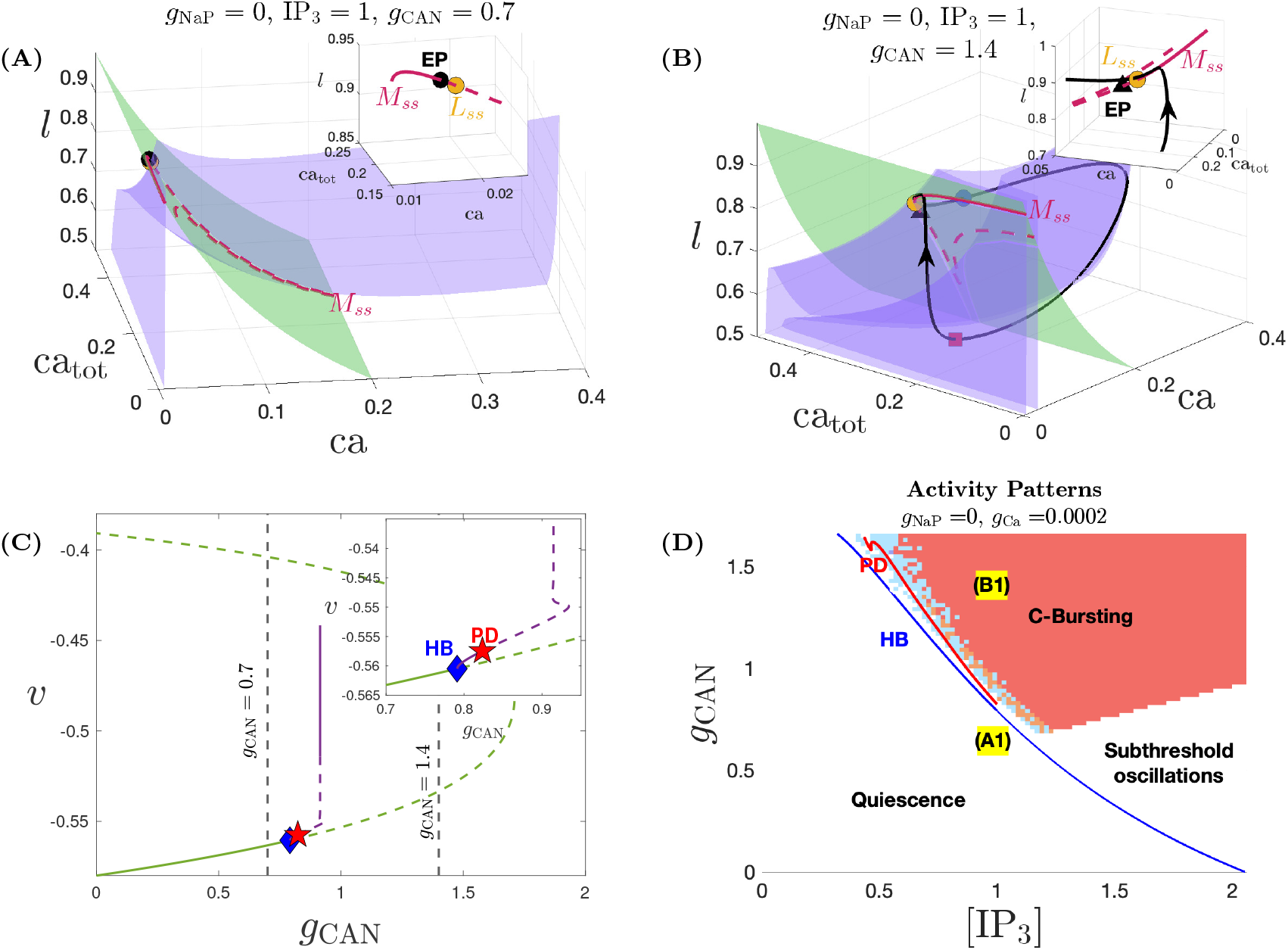
Projections of the solutions from Figure 9A1 and B1 onto (*ca, ca*_*tot*_, *l*)-space are shown in panels (A) and (B), respectively. Also shown are the *ca*-nullsfurces, *l*-nullsurface and the superslow manifold *M*_*ss*_. Full system equilibria are shown by black circle (stable) and black triangle (unstable). The yellow circle denotes the superslow manifold fold *L*_*ss*_. Other color coding and symbols are the same as in Figure 5B. (C) Bifurcation diagram of the full system (2.6) with *g*_CAN_ as a parameter, for *g*_NaP_ = 0, [IP_3_] = 1 and *g*_Ca_ = 0.0002. Green and purple curves represent equilibria and family of periodic orbit solutions. Stable and unstable objects are denoted by solid and dashed lines, respectively. Inset: transitions from quiescence to small-amplitude oscillations to bursting branch via HB (blue diamond) and period-doubling PD (red star) bifurcations. (D) Two-parameter bifurcation diagram in ([IP_3_], *g*_CAN_) plane identifying the boundaries between different patterns based on the HB and PD bifurcations from panel (C), superimposed with the activity patterns for the full system (2.6).

Figure 11C shows the bifurcation structure of the full system for *g*_NaP_ = 0 and [IP_3_] = 1, using *g*_CAN_ as the bifurcation parameter. As *g*_CAN_ increases, the equilibrium changes from stable (green solid) to unstable (green dashed) via a supercritical Hopf bifurcation (blue diamond), at which a family of limit cycles emerges (purple curve). This branch loses stability through a period-doubling (PD) bifurcation (red star), after which the system exhibits bursting dynamics. Using this one-parameter diagram, we can construct a two-parameter diagram to identify regions in parameter space in which bursting persists in the absence of *I*_NaP_, i.e., the C-bursting region. Figure 11D shows the ([IP_3_], g_CAN_)-diagram for *g*_NaP_ = 0, which consists of three main distinct dynamic regimes: quiescence, subthreshold oscillations, and C-bursting. Interestingly, mixed-mode bursting oscillations (MMBOs) were also found to coexist with small-amplitude oscillations in the region between the HB and PD curves (the light blue region). While intracellular calcium *ca* oscillations emerge above the HB curve in Figure 11D, C-bursting occurs only for relatively large *g*_CAN_ (see the red region). This is because, at low *g*_CAN_, the amplitudes of *ca* oscillations are generally too small to sufficiently depolarize the membrane voltage and induce bursting. As a result, the influence of *ca* dynamics is not effectively transmitted to the voltage compartment through *I*_CAN_ and cannot sustain bursting in *v*. Using this bifurcation analysis (Figure 11C and D), we can now understand why a combined increase in [IP_3_] and *g*_CAN_ is necessary to model NE-induced C-bursting from the tonic spiking regime associated with low [IP_3_] and *g*_CAN_ values. The *g*_CAN_ values corresponding to Figure 9(A1) and (B1) are indicated by the two vertical black lines in Figure 11C, and by points (A1) and (B1) in Figure 11D. An increase in [IP_3_] alone leads to bursting that transitions to quiescence when *g*_NaP_ is blocked (point (A1) in Figure 11D), whereas increasing both *g*_CAN_ and [IP_3_] allows the system to cross the HB bifurcation and reach the C-bursting regime that persists without *I*_NaP_ (point (B1) in Figure 11D). Although a further increase in [IP_3_] for a fixed low *g*_CAN_ can also allow the system to cross the HB bifurcation, it results in subthreshold oscillations instead of bursting in the absence of *I*_NaP_, as discussed previously.

Lastly, it is worth noting that the level of *g*_Ca_ plays a critical role in determining whether C-bursting can be induced from a tonic spiking state. For example, numerical simulations in the low-*g*_Ca_ tonic spiking regime (Figure 1(i), light blue region) reveal that when *g*_Ca_ is too low (e.g., below 0.0001), increasing *g*_CAN_ and [IP_3_] fails to induce C-bursting. Additional analysis of this phenomenon is provided in Appendix E.

### 3.4. Effect of NE on Silent neurons

Our modeling also shows that quiescent neurons - those in the white region in Figure 9(i) - can switch to (N)C-bursting upon increases in *g*_CAN_ and/or [IP_3_] (Figure 9(ii)). This outcome, however, contradicts experimental observations showing that certain silent neurons remain inactive in the presence of NE. To reconcile this discrepancy, we used our model to identify conditions under which silence is preserved despite NE exposure. Our analysis predicts that neurons with low *g*_Ca_ values (e.g., *g*_Ca_ *<* 0.0001) remain quiescent even when both *g*_CAN_ and [IP_3_] are elevated (compare Figure 12(i) and (ii), white region). In these neurons, the calcium current is too weak to support intrinsic calcium oscillations, and in the absence of sufficient *I*_NaP_, neither (N)C- nor C-bursting mechanisms can be activated (see Figure 14 for supporting analysis). Thus, our model predicts a potential mechanistic explanation for the persistence of silence in a subset of neurons exposed to NE.

**Fig. 12:**
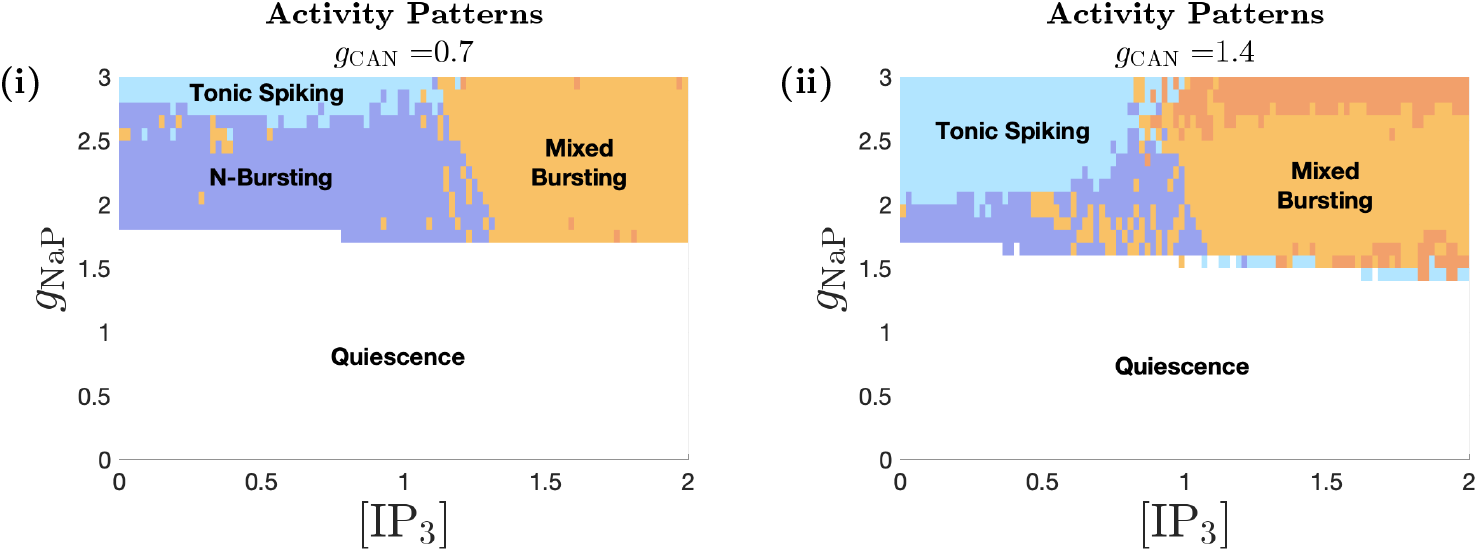
Activity patterns of model (2.6) for *g*_Ca_ = 5*e* − 5, shown as a function of *g*_NaP_ and [IP_3_], with (i) *g*_CAN_ = 0.7 and (ii) *g*_CAN_ = 1.4. Color coding follows that in Figure 1(i). In this diagram, the majority of quiescent (silent) neurons remain silent with increase in both *g*_CAN_ and [IP_3_], while other activity pattern regions either shrink or expand.

**Fig. 13:**
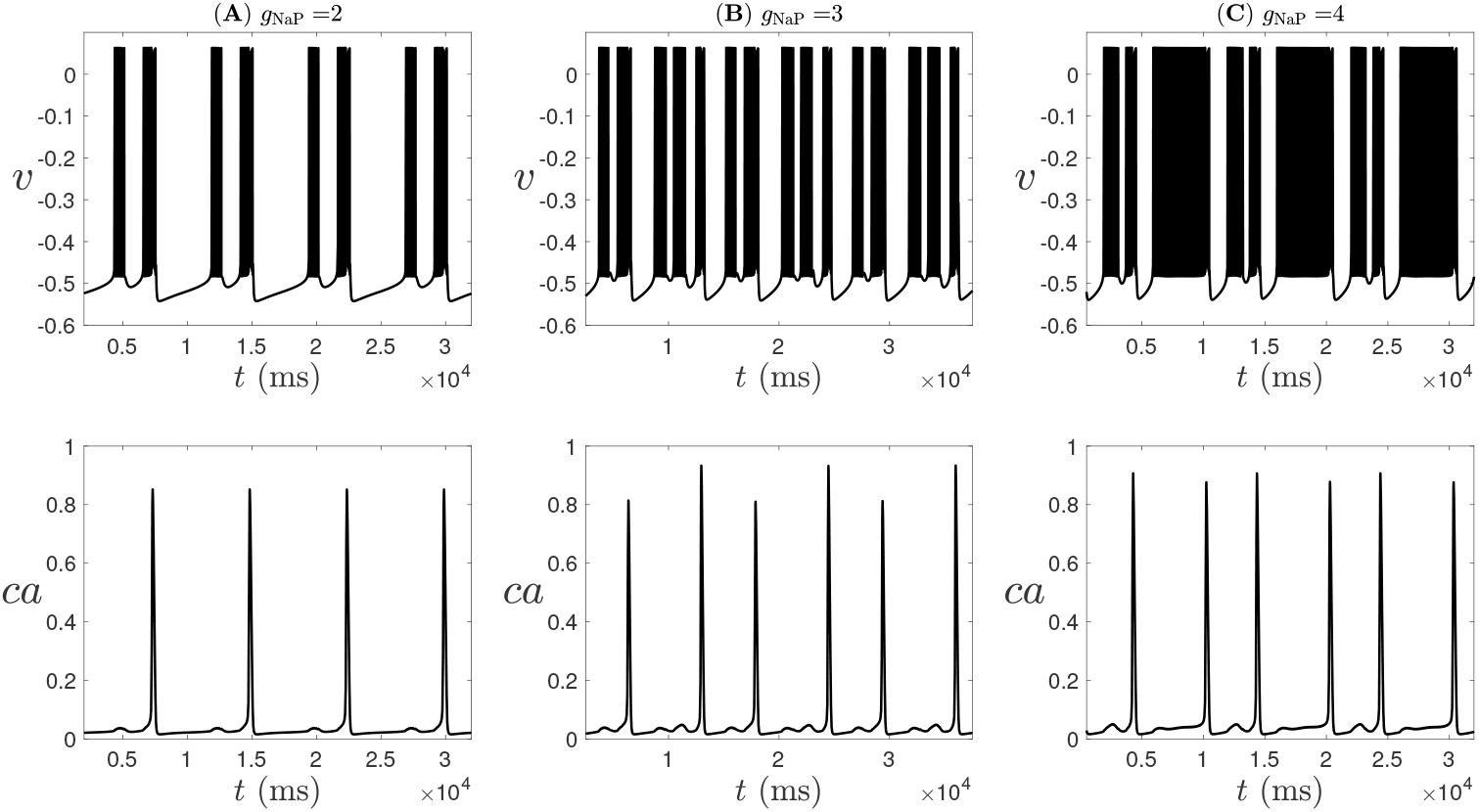
Some example voltage and calcium temporal traces from (2.6) exhibiting mixed bursting, occurring during the transition from tonic spiking to induced (N)C-bursting (see Figure 9), with parameters: *g*_Ca_ = 0.0002, *g*_CAN_ = 0.7, [IP_3_] = 0.4, and (A) *g*_NaP_ = 2, (B) *g*_NaP_ = 3, (C) *g*_NaP_ = 4.

**Fig. 14:**
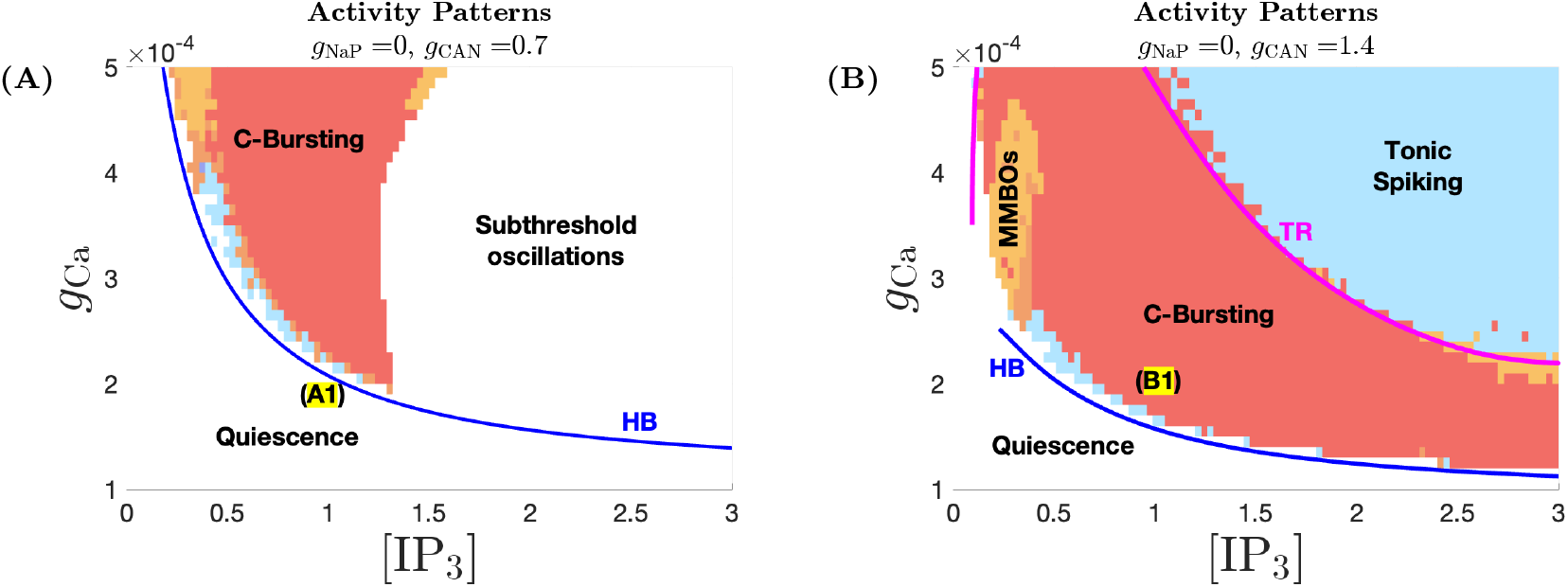
Two-parameter bifurcation diagrams of the full system (2.6) in the ([IP_3_], *g*_Ca_)-space, with *g*_NaP_ = 0. (A) *g*_CAN_ = 0.7; (B) *g*_CAN_ = 1.4. The Hopf bifurcation curve (HB) is shown in blue, and the torus bifurcation curve (TR) is shown in magenta. Activity patterns of the full system (2.6) are superimposed.

## 4. Discussion

Despite previous experimental and computational efforts in understanding the basis for the modulation of norepinephrine (NE) in the preBötzinger complex (preBötC), there has been limited progress toward understanding the mechanisms by which NE differentially impacts distinct preBötC neuronal types. Our paper addresses this gap by proposing a novel mechanism wherein NE enhances neuronal excitability through the concurrent upregulation of calcium-activated non-specific cation conductance (*g*_CAN_) and intracellular inositol trisphosphate (IP_3_) levels. This dual-modulation mechanism engages second messenger-mediated calcium dynamics and successfully captures NE-induced conditional bursting in neurons that in synaptic isolation are tonically spiking, a phenomenon not seen in earlier modeling studies that considered only NE-enhanced *g*_CAN_ [51, 38]. Our *in silico* model also identifies parameter regimes in which neurons remain quiescent despite elevated levels of *g*_CAN_ and [IP_3_]. In addition, it reveals distinct

NE-induced changes in burst frequency and duration between NaP- and CAN-dependent bursters, providing a mechanistic explanation for the experimentally observed heterogeneity in NE responsiveness. Collectively, these findings reconcile discrepancies between previous computational and experimental studies by reproducing key experimental observations and offer novel insights into the diverse modulatory roles of NE within the preBötC network.

To gain deeper insight into the proposed dual-modulation mechanism of NE, we performed GSPT analysis to analyze how this mechanism affects distinct neuronal types. This analysis predicts that conditional bursting in tonic spiking neurons requires a concurrent elevation of both [IP_3_] and *g*_CAN_. As illustrated in subsection 3.3, increasing *g*_CAN_ alone is insufficient to induce bursting; in fact, a sufficient increase in *g*_CAN_ can even switch a bursting neuron to tonic spiking, opposite to the transition induced by NE. While increasing [IP_3_] can induce bursting in some tonic spiking neurons, the resulting bursting is not solely dependent on *I*_CAN_, as eliminating either *I*_NaP_ or *I*_CAN_ abolishes the bursting dynamics. Our analysis predicts that only when both *g*_CAN_ and [IP_3_] are sufficiently elevated does the induced bursting become strictly CAN-dependent, mimicking the NE-induced bursting properties. Moreover, we demonstrate that such NE-induced bursting exhibits a distinct mechanism from that of the regular CAN-dependent burster analyzed in subsection 3.2 (compare Figures 5 and 10).

Our analysis of NaP-dependent bursting (N-bursting) neurons in subsection 3.1 demonstrates that NE increases N-burst frequency while having minimal impact on burst duration (Figure 3), consistent with experimental findings. This effect occurs primarily through the increase of *g*_CAN_, which both decreases the distance between *I*_NaP_ inactivation thresholds for burst initiation and termination (i.e., (*h*_SN_ − *h*_HC_) in Figure 2C) and accelerates the increasing rate of *h*_NaP_ during the silent phase, together leading to a shortened interburst interval and thus a higher burst frequency. In contrast, during the active phase, the effects of *g*_CAN_ on (*h*_SN_ − *h*_HC_) and the rate of *h*_NaP_ are opposing, effectively counterbalancing each other to maintain a nearly constant burst duration. Interestingly, our simulations suggest that the increase in frequency occurs in discrete steps as the model reaches certain thresholds in *g*_CAN_ values. With higher [IP_3_], these thresholds shift slightly to higher *g*_CAN_ values. Our analysis suggests that each threshold corresponds to a saddle-node bifurcation of the bursting branch in the full system (see Figure 4). Between bifurcations, the burst frequency remains nearly constant, but experiences a sudden increase upon crossing each bifurcation threshold, resulting in a phasic pattern of burst frequency in the ([IP_3_], *g*_CAN_) parameter space. This pattern of discrete changes is also reflected in the N-burst duration (see Figure 3(ii)), though we only observe a sudden increase in burst duration along the bifurcation threshold curve, while the N-burst duration otherwise remains nearly constant. Understanding the mechanisms that maintain homeostatic properties between bifurcation thresholds, and the abrupt changes across them, remain an interesting direction for future research. Based on this, we predict that the baseline conditions and the conditions where NE concentrations change correspond to different homeostatic regions in the ([IP_3_], *g*_CAN_) parameter space. Whether a similar phasic pattern also exists in experimental data remains a question, but can be tested through dose-response experiments with NE, as existing literature primarily focuses on single-dose responses.

Our analysis shows that CAN-dependent bursting (C-bursting) is driven by calcium oscillations via *I*_CAN_ activation, with burst onset mediated through a delayed HB on the superslow manifold *M*_*ss*_. As explained in subsection 3.2, increasing *g*_CAN_ alone increases burst frequency, whereas an initial increase in [IP_3_] slows down the frequency, followed by acceleration with further increases in [IP_3_]. As a result, there exist regions in the ([IP_3_], *g*_CAN_)-parameter space where the C-burst frequency remains nearly constant despite increases in both parameters. This highlights the importance of incorporating both mechanisms to reproduce the experimental observation that the C-burst frequency remains nearly unchanged under NE application. Nonetheless, this unchanged frequency is accompanied by a nearly constant burst duration (Figure 8C-G). There are also regions where increasing [IP_3_] and *g*_CAN_ leads to increases in both burst frequency and duration (Figure 8G-H). In the first case, while C-burst frequency remains similar, the lack of an increase in burst duration contradicts experimental observations from [56]. In the second case, although the increase in burst duration aligns with the experimental results, it comes at the cost of an increase in burst frequency, which again differs from experimental findings.

One possible explanation for this discrepancy is that the relatively high (20 *µ*M) dose of NE used in the experimental study by [56] may have depolarized the CAN-dependent bursters sufficiently to the point where burst frequency does not further increase with more excitability. Such a burst frequency saturation effect has been shown in C-bursters where sequentially increasing depolarizing steps of current injections initially increase burst frequency, but subsequent larger depolarizations no longer increase it [50]. Thus, it is possible that NE at 20 *µ*M quickly saturated the frequency response of the C-bursters. Another possibility is that NE could activate more than one adrenergic receptor subtype, resulting in activities not captured by our modeling.

*g*_CAN_-induced increases in C-burst frequency were not observed in the earlier models [51, 38], where *ca*-oscillations acted as an independent relaxation oscillator, unaffected by membrane voltage due to the absence of *I*_Ca_. This decoupling was a consequence of the closed-cell assumption, which excluded calcium influx from the extracellular space and limited the responsiveness of C-burst frequency (i.e., the frequency of calcium oscillations) to membrane depolarization. In contrast, our model incorporates *I*_Ca_ to relax the closed-cell assumption [31, 42, 59], leading to more complex effects of *g*_CAN_ on C-bursting dynamics through the voltage dependence of the calcium subsystem, as demonstrated by our analysis in subsection 3.2. A more detailed GSPT analysis, incorporating additional subsystems and projections, may be necessary to further clarify these effects and determine how to carefully balance *g*_CAN_ and [IP_3_] to achieve an increase in burst duration while maintaining a constant burst frequency in an open cell model. Thus, although our model does not fully reproduce all experimental observations regarding the effects of NE on C-bursters, it provides specific, testable predictions about the interplay between *g*_CAN_ and [IP_3_].

While we do not vary *g*_Ca_ when modeling NE, its baseline level varies across neuronal types and can have an important impact on the effects of NE. For example, *g*_Ca_ remains relatively low in N-bursting neurons but is much higher in C-bursting neurons (see Figure 1), which partly contributes to the different effects of NE on these two bursting dynamics as discussed above. Moreover, subsection 3.3 shows that NE induces C-bursting in tonic spiking neurons with intermediate *g*_Ca_, but not in those with low *g*_Ca_. The role of *g*_Ca_ in modeling NE-induced C-bursting is further analyzed in Appendix E. Similarly, our model predicts that the effect of NE on silent neurons also critically depends on the level of *g*_Ca_. With low *g*_Ca_, the majority of silent neurons remain quiescent in the presence of NE (see Figure 12). However, as *g*_Ca_ increases, independent calcium oscillations can emerge with NE, and this quiescence is no longer preserved in silent preBötC neurons with higher levels of *g*_Ca_. Thus, our analysis predicts that neurons with relatively low *g*_Ca_ values are capable of remaining silent despite increases in both *g*_CAN_ and [IP_3_], consistent with experimental observations. This *g*_Ca_-dependent response to the application of NE in our model raises the question of whether similar differential effects occur in preBötC neurons *in vitro*, suggesting that further experimental investigation may be needed.

Our modeling also revealed a variety of interesting respiratory bursting dynamics. In modeling the effects of NE, we observed both mixed bursting (see Figure 1D in subsection 2.2 and Appendix D) and mixed-mode bursting oscillations (see Figure 8A,B in subsection 3.2). We show that mixed bursting behavior arises during the transition from a tonic spiking neuron to bursting, suggesting that an initial application of NE might induce irregular activity before stabilizing into regular bursting with continued NE application. Since existing experiments have typically used a fixed dosage of NE, testing this hypothesis would require further experimental investigation. Beyond their biological relevance, these mixed bursting and mixed-mode bursting oscillations (MMBOs) are also of interest from a dynamical systems perspective. In particular, the small-amplitude oscillations can arise either from a delayed Hopf bifurcation (DHB) on the superslow manifold *M*_*ss*_ or through a canard mechanism associated with folded node singularities along the fold of *M*_*s*_ [15]. A key singularity known as canard-delayed-Hopf singularity naturally arises in three-timescale models such as our model (2.6), and may facilitate the coexistence and interaction of the DHB and canard mechanisms to co-modulate local oscillator behaviors [41]. Uncovering the mechanisms underlying these mixed bursting and MMBO dynamics in three-timescale settings would require a more detailed GSPT analysis, as carried out in previous studies (e.g., [6, 16, 34, 17, 57, 58, 59, 41]), and represents an intriguing direction for future research.

Beyond its single-neuron effects, neuromodulation plays a crucial role in shaping rhythmogenesis at the network level. Recent studies have shown that network sensitivity to opioid-based neuromodulation is critically determined by network configuration and the intrinsic cellular properties of neurons embedded in the network [1, 8, 12]. Our findings highlight the complex ways in which NE differentially modulates distinct neuronal subtypes, supporting the view that neurons play distinct roles in shaping network-level responses based on their intrinsic states. Extending our modeling framework to the network level will be crucial for understanding how NE-induced, cell-specific changes influence coordinated activity across the preBötC network, and how these effects interact with network architecture.

More broadly, the role of neuromodulation in rhythmogenesis is known to depend on physiological context. While NE can stabilize inspiratory activity in healthy networks [56], it may destabilize inspiratory network rhythms under pathological conditions [60]. For example, under acute intermittent hypoxia, NE has been shown to promote variability in respiratory rhythms and impair transmission to motor outputs—effects implicated in disorders such as sleep apnea and apneas of prematurity [60, 24, 7]. Although these disruptions were originally attributed to changes in inhibitory synaptic drive, our findings raise an alternative, testable hypothesis: that variability in intrinsic NE responsiveness among preBötC neurons may also contribute to such divergent outcomes. Our study lays the groundwork for future investigations into how cell-specific modulation may shape network-level effects under both healthy and disease conditions.

## Appendix A. Mathematical Preliminaries

In 1952, Hodgkin and Huxley [29] developed a mathematical model of the squid giant axon using a system of differential equations. In this paper, we model the individual preBötC neuron as a single-compartment unit with Hodgkin-Huxley conductances adapted from previously described models [38, 59]. It is well established that neural processes can evolve over very different time scales. Identifying relevant time scales is a useful step when modeling neural systems. In mathematical analysis, the time scales in a system of differential equations are often grouped into a small number of classes to facilitate simplification of analysis via limiting processes. This enables the application of a method known as *Geometric Singular Perturbation Theory* (GSPT), as we review in subsection 2.4. GSPT leverages insights from the reduced “limit systems”, which are obtained by taking the singular limit at which certain variables evolve infinitely fast or slow, to gain information about the behavior of the original system. Examples of this approach are ubiquitous in the literature, with ideas applicable to systems in which time scales can be grouped into two classes being particularly well developed. Typical simple models of membrane potential oscillations have at least two time scales. In our model, which combines membrane potential and intracellular calcium dynamics, dimensional analysis in Appendix B reveals three distinct timescales. This motivates the use of a three-timescale extension of the GSPT framework [36] to decompose the full system into fast, slow and superslow subsystems (see Appendix C for their derivations). The aim of GSPT is to combine information of these reduced systems to infer dynamics in the full systems.

In addition to GSPT, we also make extensive use of bifurcation analysis to analyze the system’s dynamic behaviors. Many interesting transitions in activity arise at bifurcation points, locations where the stability, number, and/or type of equilibria and periodic orbits change. To carry out this analysis, we construct *one-parameter bifurcation diagrams*, which track how equilibria and periodic orbits evolve as a single *bifurcation parameter* varies. These parameters may be biologically meaningful system parameters, or, in systems with multiple timescales, a slow variable can be treated as a bifurcation parameter for the fast subsystem in combination with GSPT. We use this method in our analysis of the N-bursting solution in subsection 3.1. Such bifurcation techniques have been widely used in the study of neuronal bursting dynamics [46, 5, 30, 25], including in models of respiratory neurons [38, 57, 58, 59]. Further, since biological systems are often influenced by multiple factors that can change simultaneously, we also make use of *2-parameter bifurcation diagrams*, which track how bifurcation curves change as a function of two parameters. We apply this analysis in Sections 3.1 and 3.3 to study our model dynamics, with examples shown in Figures 2C and 11D.

In Appendices B and C below, we detail other mathematical procedures such as nondimensionalization of the model and preliminaries of GSPT necessary for the analysis of our model.

## Appendix B. Nondimensionalization of the Model

In order to verify whether our model evolves in multiple timescales, we need to nondimensionalize our model. As a first step towards that, we rescale the variables so that the different timescales can be clearly identified, and to do so, we define new dimensionless variables *τ, v, ca, ca*_*tot*_ such that

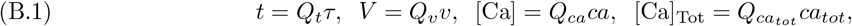

where *Q*_*t*_, *Q*_*v*_, *Q*_*ca*_ and 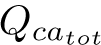 are time, voltage, calcium, and total intracellular calcium scales, respectively. We note that *n, h*, and *l* are already dimensionless variables.

Next, from numerical simulations, we find that the membrane potential *V* lies between -60 mV and 10 mV. Correspondingly, we define 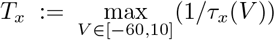, for *x* ∈ {*n, h*}. Then, define *t*_*x*_(*V*) := *T*_*x*_*τ*_*x*_(*V*), a rescaled version of *τ*_*x*_. We also define *g*_max_ := max{*g*_L_, *g*_K_, *g*_Na_, *g*_NaP_, *g*_CAN_, *g*_Ca_}. Further, we let 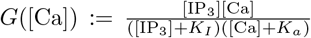 and 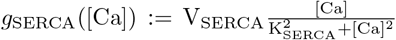. Then, 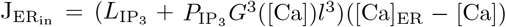 and 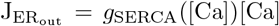. Substituting these in equation (2.1) and rearranging, we get the following system:

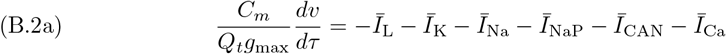

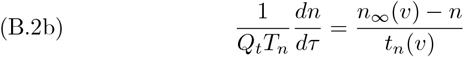

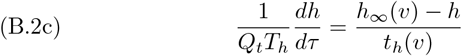

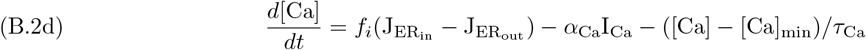

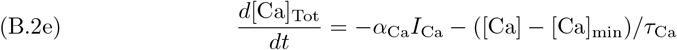

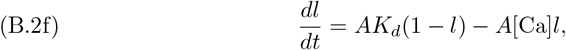

with dimensionless currents 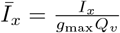.

Thus, we have nondimensionalized the voltage subsystem (B.2a) - (B.2c). Next, we focus on the calcium subsystem (B.2d) - (B.2f). Again, from numerical simulations, we know that [Ca] ∈ [0, 2] *µ*M and [Ca]_Tot_ ∈ [0, 5] *µ*M. Therefore, we define 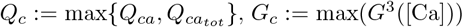 and *G*_*S*_ := max(*g*_SERCA_([Ca])) over the range [Ca] ∈ [0, 2]. Then, define 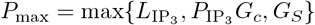. Making the necessary substitutions in and reducing the equations from (B.2), we then obtain the following system of equations:

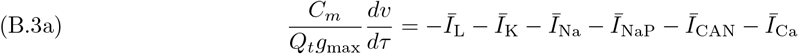

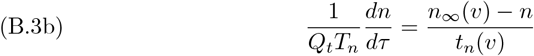

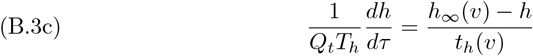

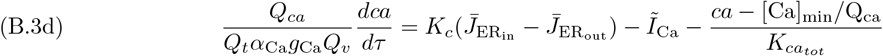

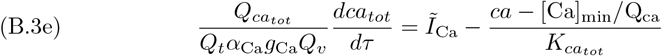

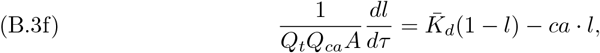

where 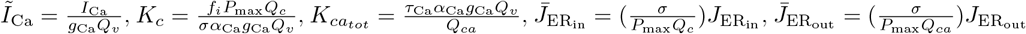 and 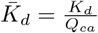.

From expected ranges for *V*, [Ca] and [Ca]_Tot_ mentioned a priori, suitable choices for the voltage and calcium scales are *Q*_*v*_ = 100 mV, *Q*_*ca*_ = 2 *µ*M and 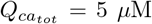, respectively, which also then implies that *Q*_*c*_ = 5 *µ*M. We note that the values of *n*_∞_, *h*_∞_, *n, h, l* ∈ [0, 1]. Moreover, for the values specified in Table 1, the maximum of the conductances is *g*_Na_ = 28 nS. Further, from numerical evaluations of 1*/τ*_*n*_(*V*) and 1*/τ*_*h*_(*V*), for *V* ∈ [−60, 10], we find that *T*_*n*_ ≈ 6.5491 ms^−1^ and *T*_*h*_ ≈ 0.0165 ms^−1^. We also compute *G*_*c*_ ≈ 0.0214 and *G*_*S*_ ≈ 1000 pL · ms^−1^, so we then have that *P*_max_ ≈ 1000 pL · ms^−1^. Since we vary *g*_Ca_ ∈ [0.00002, 0.0008] in our model, we find that this leads to *K*_*c*_ ≈ 12416 = *O*(10^4^) and 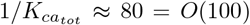 for *g*_Ca_ *<* 0.0001, while *K*_*c*_ ≈ 1241 = *O*(1000) and 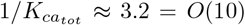 for 0.0001 ≤ *g*_Ca_ ≤ 0.0008.

Using these values, we observe that all quantities on the right-hand side of equations (B.3a) - (B.3f) except *K*_*c*_ and 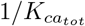 are bounded by 1 in absolute values. To resolve this issue, we then divide both sides of equation (B.3d) by *K*_*c*_ and multiply both sides of equation (B.3e) by 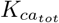. Then, we have the following set of equations:

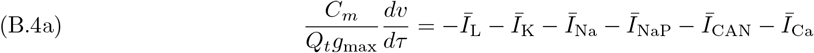

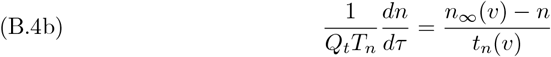

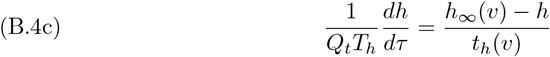

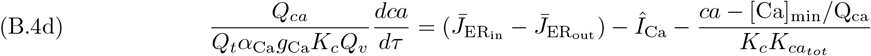

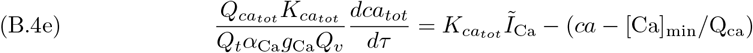

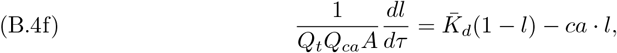

where 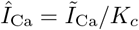.

Now, all equations on the right-hand side of (B.4) are bounded above by 1 in absolute values. Since *K*_c_ ≫1, the term 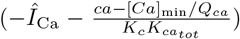 is negligible. Hence, the coefficients of the derivatives on the left-hand side of all the equations in (B.4) convey the relative rates of evolution of the variables. Since 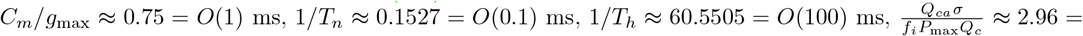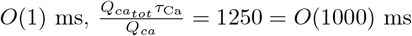 and 1*/*(*Q*_*ca*_*A*) = 100 = *O*(100) ms, we can conclude that *v, n* and *ca* evolve on a relatively fast timescale, *h* and *l* evolve on a slow timescale, while *ca*_*tot*_ evolves on a superslow timescale. We choose the slow timescale as our reference time by letting *Q*_*t*_ = 100 ms, and set:

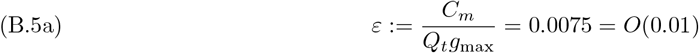

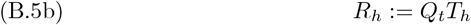

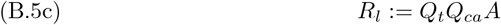

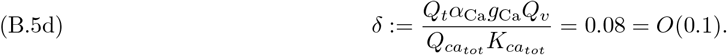

Substituting the above expressions (B.5) in (B.4) and rearranging necessary terms, we obtain:

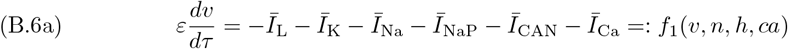

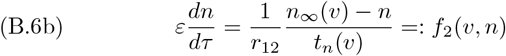

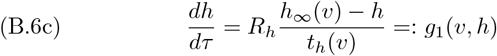

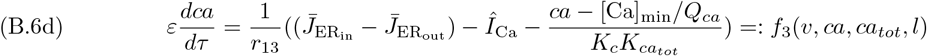

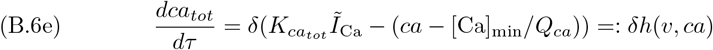

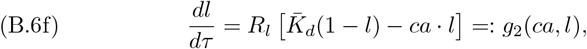

where *ε, δ* ≪ 1,

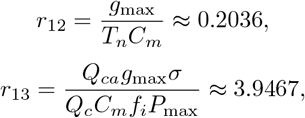

*R*_*h*_ and *R*_*l*_ are dimensionless parameters, and all functions on the right-hand side of (B.6a) - (B.6f) are bounded by 1 in absolute value and are *O*(1).

## Appendix C. Singular limits in the Nondimensionalized Model

In this section, we perform GSPT analysis on (B.6) by treating *ε* and *δ* as two independent singular perturbation parameters. Since the primary focus of this work is to explore the effects of norepinephrine on the preBötC neuron, we only provide a brief overview of the GSPT analysis and the derivation of key subsystems for the sake of completeness. Readers interested in the full details are referred to [36, 41].

Choose *τ* = *t*_*s*_ to be the reference time, we call the dimensionless system (B.6) that evolves over the *slow time t*_*s*_ the *slow system*. Introducing a superslow time *t*_*ss*_ = *δt*_*s*_ yields the following equivalent descriptions of dynamics:

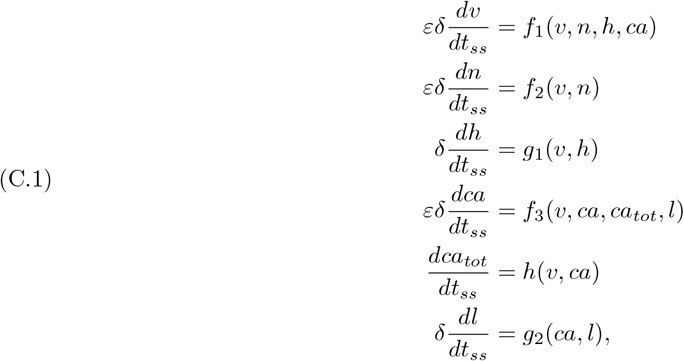

which evolves on the *superslow timescale* and is called the *superslow system*. Alternatively, defining a fast time *t*_*f*_ = *t*_*s*_*/ε* leads to another equivalent *fast system*:

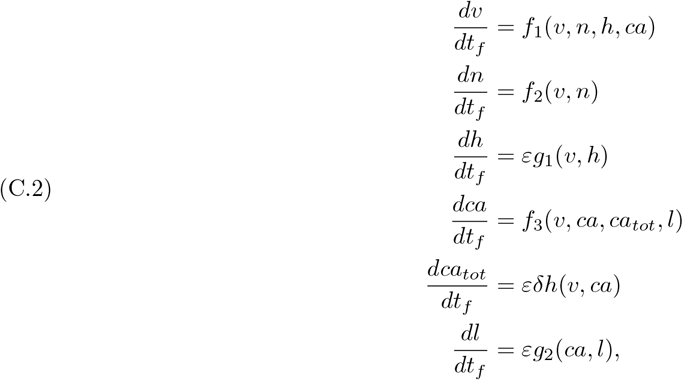

which evolves on the *fast timescale*.

The existence of two independent singular perturbation parameters, *ε* and *δ*, implies there are various ways to implement GSPT, each yielding distinct singular limit predictions.

### Singular Limit as *ε* → 0

Fixing *δ >* 0 and letting *ε* → 0 in the fast system (C.2) yields the three-dimenstional (3D) *fast layer problem*, a system that describes the dynamics of the fast variables *v, n* and *ca* for fixed values of *h, l* and *ca*_*tot*_:

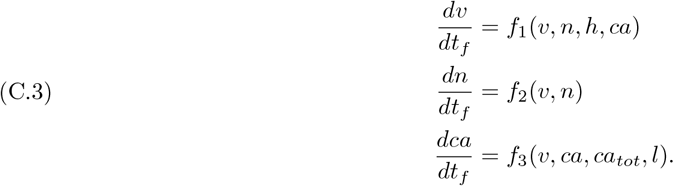

The set of equilibrium points of the *fast layer problem* is called the *critical manifold*, denoted by *M*_*s*_:

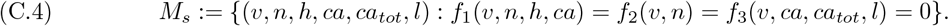

For sufficiently small *ε >* 0, normally hyperbolic parts of *M*_*s*_ perturb to a locally invariant manifold called a *slow manifold* [22]. In this paper, for simplicity, we simply use *M*_*s*_ as a convenient numerical approximation of these slow manifolds.

*M*_*s*_ is folded with a set of saddle-node bifurcations of the fast subsystem (C.5) *L*_*s*_ := {(*v, n, h, ca, ca*_*tot*_, *l*) ∈ *M*_*s*_ : det(*J*) = 0},

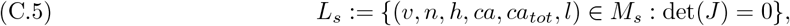

where det(*J*) denotes the determinant of the 3 × 3 Jacobian matrix of the fast subsystem (C.3). The fold curves *L*_*s*_ separate the attracting and repelling branches of the critical manifold *M*_*s*_.

Taking the same limit *ε* → 0 with *δ >* 0 in the slow system (2.6) yields the 3D *slow reduced problem*, which describes the dynamics of the slow and the superslow variables along the critical manifold *M*_*s*_ (2.7):

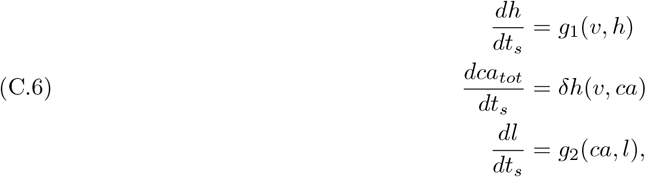

*where f*_1_ = *f*_2_ = *f*_3_ = 0.

### Singular Limit as *δ* → 0

Alternatively, letting *δ* → 0 and fixing *ε >* 0 in the slow system (2.6) yields the 5D *slow layer problem*:

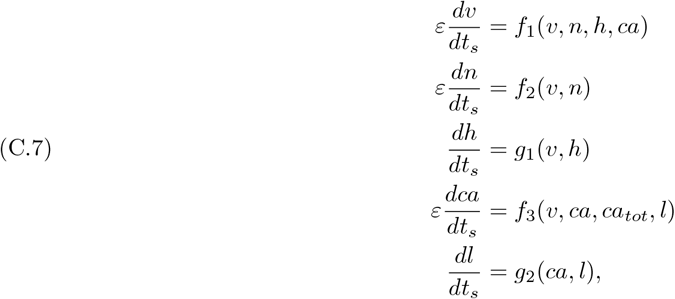

where the superslow variable *ca*_*tot*_ is a constant parameter.

The set of equilibrium points of the slow layer problem (C.7) is a one-dimensional subset of *M*_*s*_ called the *superslow manifold* and is denoted by *M*_*ss*_:

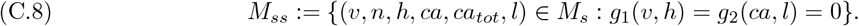

Again, similar to *M*_*s*_, for 0 *< δ* ≪ 1, the normally hyperbolic parts of *M*_*ss*_ perturb to locally invariant manifolds, which we shall simply refer to as *M*_*ss*_. Interesting dynamics are expected to occur near nonhyperbolic points on *M*_*ss*_ where Fenichel’s theory (GSPT) breaks down. These include Hopf or saddle-node fold bifurcations of the slow layer problem (C.7).

Taking the same limit, i.e., *δ* → 0 with *ε >* 0, in the superslow system (C.1) leads to the *superslow reduced problem* representing the dynamics of the superslow variable *ca*_*tot*_ along the *M*_*ss*_ (2.8):

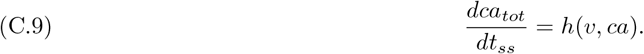

where *f*_1_ = *f*_2_ = *f*_3_ = *g*_1_ = *g*_2_ = 0. The superslow dynamics of (C.9) are slaved to *M*_*ss*_ until nonhyperbolic points are reached.

### Double Singular Limits as *ε* → 0 and *δ* → 0

Further, since both the slow reduced problem (C.6) and the slow layer problem (C.7) still evolve on two distinct timescales, taking the limit *δ* → 0 in (C.6) or *ε* → 0 in (C.7) yields the same *slow reduced layer problem*:

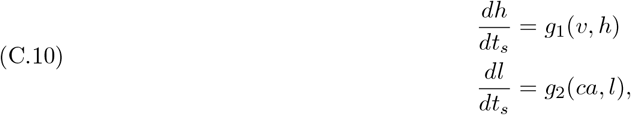

which describes the dynamics of the slow variables *h* and *l* along *M*_*s*_ and the superslow variable *ca*_*tot*_ is a constant.

## Appendix D. Mixed Bursting Dynamics

In subsection 3.3, when a tonic spiking neuron switches to a bursting behavior, there exists an intermediate parameter region along the transitional region in the ([IP_3_], *g*_NaP_)-space (see Figure 9(i)) where the model exhibits some interesting *mixed bursting* dynamics. We use the term mixed bursting to describe all such dynamics where a period may include two or more bursts, with each of these bursts having similar or different burst widths. These mixed bursts persist with an increase in *g*_CAN_ (Figure 9(ii)) as well as with small variations in *g*_Ca_ (Figure 1(i)).

In our model, this mixed bursting behavior is a subclass of Type 1 (N)C-bursters in which the bursting mechanism depends on both the NaP and CAN mechanisms in such a way that in the absence of either, the bursts are lost. Such a mixed bursting behavior arises for *g*_Ca_ ≈ 0.0002, which is just sufficient to produce independent *ca*-oscillations. These kinds of bursts are especially interesting since, from Figure 13, looking at the *ca*-dynamics, we note multiple small-amplitude oscillations that occur before a full oscillation in the *ca*-subsystem. This suggests the presence of canard-like dynamics in our model. Moreover, the fact that such bursts occur over a relatively large range of *g*_NaP_, as well as *g*_Ca_ and [IP_3_] to a reasonable extent, makes them a compelling behavior in need of further analytical exploration.

These bursts are also interesting from the perspective of understanding the effect of NE in our model. Because they occur for an intermediate range of parameter values observed during the transition of a tonic spiking neuron to an induced (N)C-burster, this might suggest that the concentration of NE applied may play an important role in seeing a successful transition. Further, the concentration could also be a factor of consideration at the network level in studying the network synchrony.

## Appendix E. Role of *g*_Ca_ in modeling NE-induced C-bursting

Recall from subsection 3.3 that in model (2.6), tonic spiking arises under two distinct parameter regimes: when calcium conductance (*g*_Ca_) is low (less than 0.0001) (see Figure 1(i), light blue region), or when *g*_Ca_ is intermediate (e.g., ∼ 0.0002) (see Figure 9(i), light blue region). In both cases, *g*_NaP_ needs to be relatively high to sustain tonic spiking in the voltage dynamics. As shown in subsection 3.3, while NE can induce C-bursting from the tonic spiking regime with intermediate *g*_Ca_, it fails to do so when *g*_Ca_ is too low.

To understand this, we compute the bifurcation structure of the full system (2.6) with respect to [IP_3_] and *g*_Ca_, again setting *g*_NaP_ = 0 (see Figure 14). It is important to note that only the quiescent region with low or intermediate *g*_Ca_ and low [IP_3_] in Figure 14 corresponds to the tonic spiking regime in the full system when *g*_NaP_ is high. To examine the condition on *g*_Ca_ under which C-bursting can be induced, we restrict our attention to the range *g*_Ca_ ≤ 0.0002, ensuring the system begins in a tonic spiking state at high *g*_NaP_ and low [IP_3_]. Figure 14 shows that increasing *g*_CAN_ from 0.7 (panel (A)) to 1.4 (panel (B)) shifts the HB curve to lower *g*_Ca_, thereby expanding the C-bursting region. Moreover, subthreshold oscillations above the HB curve are eliminated with increased *g*_CAN_, while tonic spiking solutions are observed in the light blue region above the TR bifurcation curve (magenta). It follows that when *g*_Ca_ is below the horizontal asymptote of the HB curve at high *g*_CAN_ (e.g., *g*_Ca_ *<* 0.00012), increasing *g*_CAN_ and [IP_3_] is insufficient to cross the HB curve and induce a transition to C-bursting. In contrast, for intermediate *g*_Ca_ (e.g., *g*_Ca_ = 0.0002 at points (A1) and (B1)), increasing *g*_CAN_ and [IP_3_] can drive a neuron —initially silent in the absence of *g*_NaP_ but corresponding to tonic spiking at high *g*_Nap_— across the HB bifurcation and transition to a C-bursting state that persists without *I*_NaP_.

## Acknowledgments

We thank Dr. Jean-Charles Viemari for helpful comments on this work. This work was supported by NIH/NIDA R01DA057767, as part of the Collaborative Research in Computational Neuroscience Program. A.J. Garcia 3rd is also supported by NIH/NHLBI R01HL163965 and NIH/NIDA R01DA061412.

